# RADX Sustains POT1 Function at Telomeres to Counteract RAD51 Binding, which Triggers Telomere Fragility

**DOI:** 10.1101/2020.01.20.912634

**Authors:** Anna-Sophia Briod, Galina Glousker, Joachim Lingner

## Abstract

The 3’ terminal DNA extensions at chromosome ends can become engaged into multiple biochemical reactions during DNA replication, telomerase-mediated telomere extension, homology directed DNA repair, nucleolytic processing and DNA damage checkpoint activation. To keep these activities in check, telomeric 3’ overhangs can be hidden in t-loop structures or they associate with specialized proteins such as POT1. Here, we explore the telomeric microenvironment using a proximity-dependent labeling approach and identify the oligonucleotide/oligosaccharide-binding (OB)-fold containing protein RADX. RADX binds single-stranded telomeric DNA throughout the cell cycle along with POT1, suppressing accumulation of fragile telomeres, which are indicative of telomere replication defects. Telomere fragility in POT1 and RADX double-depleted cells was due to accumulation of the RAD51 recombinase at telomeres. RADX also supports DNA damage signaling at POT1-depleted telomeres counteracting RAD51 binding. Thus, RADX represents next to POT1 a second OB-fold containing single-strand telomere binding protein sustaining telomere protection.

## Introduction

Intact telomeres suppress at chromosome ends DNA repair activities, nucleolytic degradation and DNA damage checkpoint activation (de Lange, 2018; Lazzerini-Denchi and Sfeir, 2016). In humans, unharmed telomeres have a length of 5,000-15,000 bp of 5’-TTAGGG-3’/5’-CCCTAA-3’ telomeric DNA repeats ending in a single stranded 3’ overhang of 50-400 nucleotides. Telomeres associate with the shelterin proteins consisting of TRF1 and TRF2 which bind as homodimers to double-stranded telomeric DNA (de Lange, 2018) and POT1 which binds to the single-stranded 5’-TTAGGG-3’ repeats (Baumann and Cech, 2001). The shelterin components TIN2, RAP1 and TPP1 are recruited to telomeres through protein-protein interactions with TRF1, TRF2 and POT1. In addition to shelterin components, which are abundant at telomeres presumably covering large parts of telomeric DNA, additional factors have been identified at chromosome ends through genetic, molecular biological and biochemical approaches. More recently telomeric protein composition has been analyzed more comprehensively through the purification of crosslinked telomeric chromatin and analysis by mass spectrometry (Bartocci et al., 2014; Déjardin and Kingston, 2009; Grolimund et al., 2013; Majerska et al., 2018) or through mass spectrometric analyses of proteins that were labeled at telomeres with biotin upon expression of shelterin components fused with a biotin ligase (Garcia-Exposito et al., 2016, this study). More than 200 proteins have been identified in these studies and for a subset of them crucial functions have already been documented.

The non-shelterin telomere associated proteins become particularly important during telomere replication or telomere damage. For example, short telomeres change their state during cellular senescence triggering ATM and ATR recruitment and activation to promote permanent cell cycle arrest (d’Adda di Fagagna et al., 2003; Karlseder et al., 1999, 2002). For ATR activation, POT1 is replaced on the single-stranded telomeric DNA by RPA which recruits ATR-ATRIP (Denchi and de Lange, 2007; Zou and Elledge, 2003). Also during semiconservative DNA replication, POT1 is thought to be replaced by RPA (Flynn et al., 2011) which stimulates DNA polymerases. For telomerase-mediated telomere extension in S-phase of the cell cycle (Schmidt and Cech, 2015), telomerase engages with the telomeric 3’ overhang upon recruitment by TIN2 associated TPP1 (Abreu et al., 2010; Schmidt and Cech, 2015) and the ATM and ATR kinase activate the extension process (Lee et al., 2015; Tong et al., 2015). The OB-fold containing CST complex associates with the extended telomeric 3’ overhang to terminate telomerase-mediated telomere extension promoting fill-in synthesis of the complementary strand (Chen et al., 2012). Finally, though excessive homologous recombination between telomeres is suppressed in mouse embryonic fibroblasts (MEFs) by contributions of Pot1, Ku and Rap1 (Celli et al., 2006; Palm et al., 2009; Sfeir et al., 2010; Wu et al., 2006), the telomeric 3’ overhang can associate with RAD51 and the homology-directed DNA repair (HDR) machinery may contribute to telomere maintenance even in healthy cells (Badie et al., 2010; Verdun and Karlseder, 2006). The HDR involvement in telomere maintenance is most pronounced in ALT cells that maintain their telomeres independently of telomerase (Pickett and Reddel, 2015).

During S phase for the faithful replication of telomeric DNA, not only the canonical replication machinery must associate with telomeres but additional specialized proteins are recruited to overcome telomere-specific hurdles. For example, the t-loops are unwound by RTEL1 (Sarek et al., 2016; Vannier et al., 2012), G-quadruplex structures which can be formed during replication by the G-rich telomeric strand are counteracted by BLM, RTEL1 and WRN helicases (Crabbe et al., 2004; Sfeir et al., 2009; Vannier et al., 2012). Telomeric R-loops formed between the telomeric long noncoding RNA TERRA and the C-rich telomeric DNA strand are repressed by RNA surveillance factors (Azzalin et al., 2007; Chawla et al., 2011), RNase H (Arora et al., 2014; Graf et al., 2017), FANCM (Silva et al., 2019) and the THO-complex (Pfeiffer et al., 2013). Telomere replication defects become apparent as so-called fragile telomeres which display discontinuities of the telomeric signals in metaphase chromosomes (Sfeir et al., 2009).

Recently, the OB-fold containing protein RADX has been identified to control replication fork protection by antagonizing RAD51 binding to single stranded DNA (Bhat et al., 2018; Dungrawala et al., 2017; Schubert et al., 2017). In this paper, we explore the telomere protein composition by proximity-dependent labeling using TRF1, TRF2 and POT1 as baits. Apart from obtaining comprehensive insights into the telomeric microenvironment we identify RADX with all baits. We demonstrate that RADX associates with telomeres in replicating and non-replicating cells through its DNA binding OB-fold. Concomitant loss of RADX and POT1 leads to RAD51 recruitment and enhanced telomere fragility, which can be suppressed by RAD51 depletion. Thus, RADX contributes to telomere protection in conjunction with POT1.

## Results

### BioID Identifies RADX at Telomeres

We employed BioID (Roux et al., 2012) to explore and compare the proteomic microenvironments of TRF1, TRF2 and POT1. In order to express the biotin protein ligase (BirA) tagged shelterin proteins BirA-TRF1, BirA-TRF2 and BirA-POT1 at endogenous levels and to avoid biotinylation artifacts due to overexpression, we used a CRISPR-Cas9 knock-in approach to integrate the BirA sequence into the genomic loci at the N-termini of *TRF1*, *TRF2* and *POT1* in HEK293T cells (Figure 1A). We screened for recombinant clones by PCR with primers flanking the region of integration and sequenced the PCR products (Figure S1A, Figure S1B, Figure S1E and Figure S1G). Expression of the tagged fusion proteins was confirmed on Western blots (Figure S1C, Figure S1F, Figure S1H). Sequencing of the genotyping PCR revealed one tagged and two disrupted alleles for the myc-BirA-TRF1 clone (Figure 1A and Figure S1D), as well as for 3xFlag-BirA-POT1 clone 5 (Figure S1G and Figure S1I). Clone 3xFlag-POT1 84 showed evidence of two unedited alleles (Figure S1I) retaining expression of untagged POT1 (Figure S1H). Of note, HEK293T cells are triploid for chromosomes 7 and 8, which contain the *POT1* and *TRF1* genes, respectively (Figure 1A). The genotyping PCR for clone 3xFlag-BirA-TRF2 88 revealed the tagged allele of *TRF2* and not the unmodified locus (Figure S1E), but the Western blot analysis indicated co-expression of tagged and untagged TRF2 (Figure S1F).

**Figure 1.**
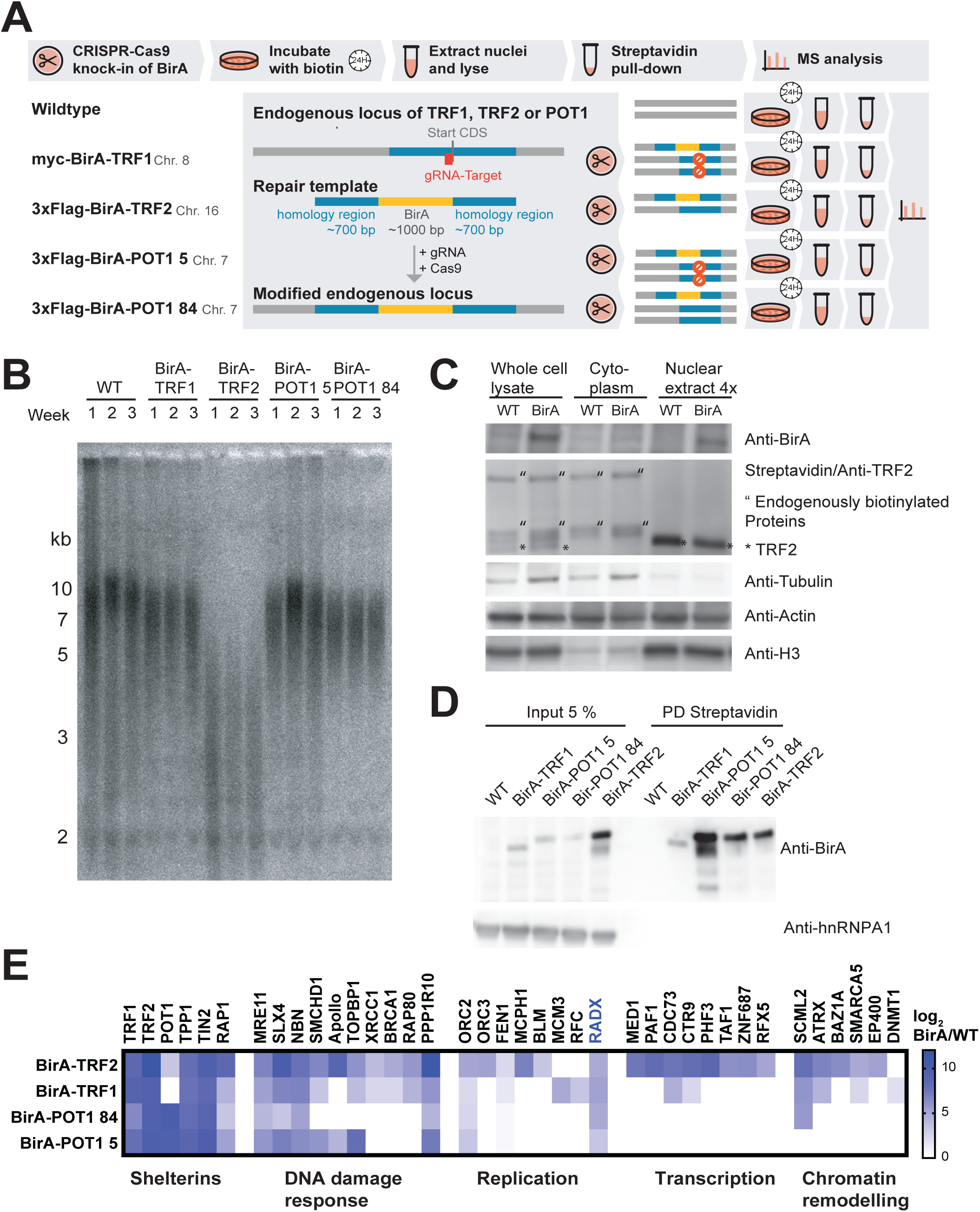
BioID identifies RADX at telomeres. (A) Schematic explaining the CRISPR/Cas9 approach to integrate the BirA sequence into the endogenous loci of TRF1, TRF2 or POT1 in HEK293T cells. Genotypes of selected clones are shown with either intact WT allele, integrated BirA or disrupted WT allele (stop sign). (B) TRF analysis of indicated clones followed over several weeks in culture. (C) Western blot showing the fractionation of endogenously biotinylated proteins present in WT and the BirA-tagged TRF1 clone 21. (D) Immunoprecipitation of BirA-tagged proteins after incubation with Biotin for 24h. (E) Heatmap showing enrichment ratios, calculated by log2 total spectral counts BirA/WT, for a selection of proteins identified in the corresponding BioID experiments. Proteins were not identified in the WT control IP or at least 3-fold enriched (total spectral counts) in BirA-IP. All proteins in this table, except for RADX had been previously identified at telomeres.

The telomere length of genome-edited cells was clone-specific as typically seen in clonal isolates of human cells and it remained stable during several weeks of growth when assessed by telomere restriction fragment (TRF) analysis indicating normal shelterin function (Figure 1B). Telomere integrity was also analyzed by inspecting telomeric fluorescence in situ hybridization (FISH) signals on metaphase chromosome spreads (Figure S2). Telomere abnormalities did not increase in the analyzed clones. Thus, the tagged proteins showed no interference with telomere maintenance.

Wild type (WT) and genome-edited cells were expanded and labeled during 24 hours with biotin. Nuclei were isolated removing cytoplasm and mitochondria, which contain abundant endogenous biotinylated proteins (Figure 1C). Biotinylated proteins in nuclear extracts were bound to streptavidin beads and fractionated by SDS-PAGE. As expected, the streptavidin-purified fractions contained also the BirA-tagged shelterins due to self-biotinylation (Figure 1D). Analysis of the streptavidin-purified and SDS-PAGE fractionated proteins by mass spectrometry and comparison to the wild type negative control led to the identification of proteins previously described to associate with telomeres (Déjardin and Kingston, 2009; Garcia-Exposito et al., 2016; Grolimund et al., 2013) as well as novel telomeric proteins (Figure 1E, S3 and Table S1). This included all the shelterin components, proteins involved in the DNA damage response, chromatin remodeling, transcription and nuclear envelope components. POT1 was not detected in the TRF1-BioID, but TRF1 in the POT1-BioID suggesting that the part of POT1 within the biotinylation radius of BirA-TRF1 does not contain primary amines accessible for biotinylation and subsequent pull-down. The overlap of with BirA-TRF1, -TRF2 and -POT1 identified proteins was extensive giving confidence in the specificity of the experimental approach although several proteins involved in transcription regulation and chromatin remodeling were only identified BirA-TRF2. However, this clone also carried shorter telomeres (Figure 1B), which may have influenced the telomeric proteome environment. Overall, we were most intrigued by the identification of RADX in all BioID experiments (Figure 1E), which had not been found in previous proteomic analyses of telomeres.

### RADX Binds Single-stranded Telomeric DNA in Interphase and S phase

RADX contains three OB-folds (Figure 2A). In previous work, OB2 was identified to be responsible for the binding of single-stranded DNA and the DNA binding activity was abrogated upon mutation of two amino acids (Dungrawala et al., 2017; Schubert et al., 2017) (Figure 2A). To test the mode of RADX association with telomeres, we transiently expressed wild type Flag-tagged RADX (3xFlagRADX) and the DNA binding defective version 3xFlag-RADX OB* in Hela cells (Figure 2A and 2B) and analyzed recruitment to telomeres by immunofluorescence (IF)-FISH. Upon detergent pre-extraction, we detected chromatin-associated 3xFlagRADX in nuclear foci, which to a large extent colocalized with telomeres (Figure 2B and 2C). Very strikingly, the size of foci, the chromatin association and telomere colocalization was strongly diminished when expressing 3xFlag-RADX OB* (Figure 2B and 2C). Consequently, we conclude that RADX associates with telomeres in dependency of its ability to bind single-stranded DNA.

**Figure 2.**
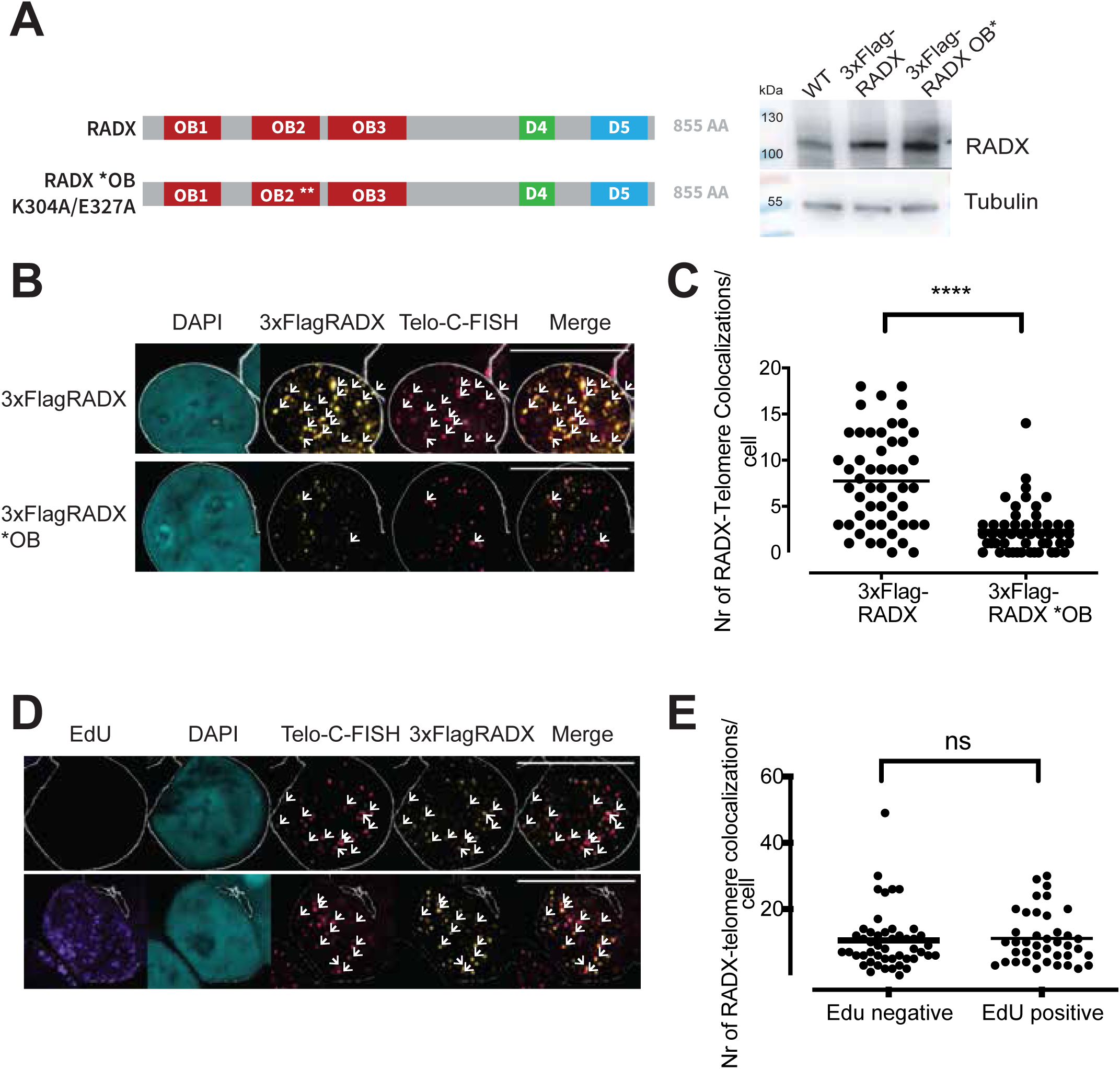
RADX binds single-stranded telomeric DNA throughout the cell cycle. (A)Schematic depicting RADX and RADX*OB with localization of the OB-folds, independent domains D4 and D5 as well as mutations in OB2. Protein levels upon transient transfection in Hela were monitored by Western Blot. (B) Transiently expressed 3XFlag-RADX (yellow) colocalizes with telomeres (pink), as indicated by white arrows. Scale bar is 10 μm. (C) Quantification of the number of RADX-telomere colocalizations per cell. At least 30 cells from three independent IF-FISH experiments were analyzed. The mean is displayed and statistical significance was calculated by unpaired t-test with p<0.0001. (D) Representative examples of IF-FISH images showing colocalization of 3XFlag-RADX with telomeres. Cells in S-Phase were determined by incubation with EdU for 10 min and labeling with a Click-it EdU Kit. (E) Quantification of number of RADX-telomere colocalizations per cell. At least 40 cells from two independent IF-FISH experiments were analyzed, the mean is displayed and statistical significance was calculated by unpaired t-test.

Since RADX was previously described to function at replication forks, we wanted to test if RADX binding to telomeres is more pronounced in S-phase (Dungrawala et al., 2017; Schubert et al., 2017). To distinguish S-phase from interphase cells, we pulse-labeled cells with the thymidine analog EdU which was subsequently fluorescently labeled with a click reaction. We observed that 3xFlagRADX colocalized with telomeres in S-phase and non-S-phase cells to a similar extent (Figure 2D and 2E). This therefore indicates that RADX associates with telomeres in interphase as well as in S-phase. Together, our experiments indicate that RADX binds to the single stranded telomeric G-rich strand which is present at the 3’ overhang of telomeres or present in the displaced G-rich strand which forms when telomeres adopt a t-loop configuration.

### HU Treatment Increases RADX Binding to Telomeres

We next sought to identify conditions which modulate RADX binding to telomeres. We tested by RNA interference (Figure 3A), if depletion of the shelterin components TRF1, TRF2 or POT1 or the RAD51 recombinase influence RADX association with telomeres. RAD51 was previously shown to antagonize RADX binding to stalled replication forks (Dungrawala et al., 2017). In addition, we tested if induction of replication stress induced by hydroxyurea (HU) or induction of DNA double-strand breaks upon zeocin treatment would influence RADX binding to telomeres. Telomere binding by RADX was assessed by chromatin immunoprecipitation (ChIP) using RADX antibodies. As a control for specificity, RADX was deleted using a specific gRNA and CRISPR/Cas9 on the population level. Telomeric DNA was detected by dotblot hybridization and Alu repeat probes were used to compare binding to a chromosome internal locus. The dotblot signal for telomeric DNA was abolished upon *RADX* deletion. RADX binding at telomeres was detected in control cells and it did not change significantly upon depletion of TRF1, TRF2 or POT1 (Figure 3B and 3C). Of note, the depletions of the shelterins were efficient as they impaired telomere function leading to increased presence of γH2AX at telomeres (Figure 3B and 3D). Zeocin treatment and RAD51 depletion increased RADX binding slightly. The most drastic increase of RADX recruitment to telomeres was observed upon HU treatment (Figure 3B and 3C). Significantly, the increased binding was observed only for the telomeric DNA but not the Alu-repeats, even though DNA damage was observed at both loci upon HU or zeocin treatment (Figure 3B and 3D). We also tested in ChIP experiments if depletion of RADX would influence POT1 binding to telomeres but did not observe an effect (Figure S4). Altogether, the results indicated that replication stress and DNA damage stimulate RADX binding at telomeres and that RAD51 counteracts this association.

**Figure 3.**
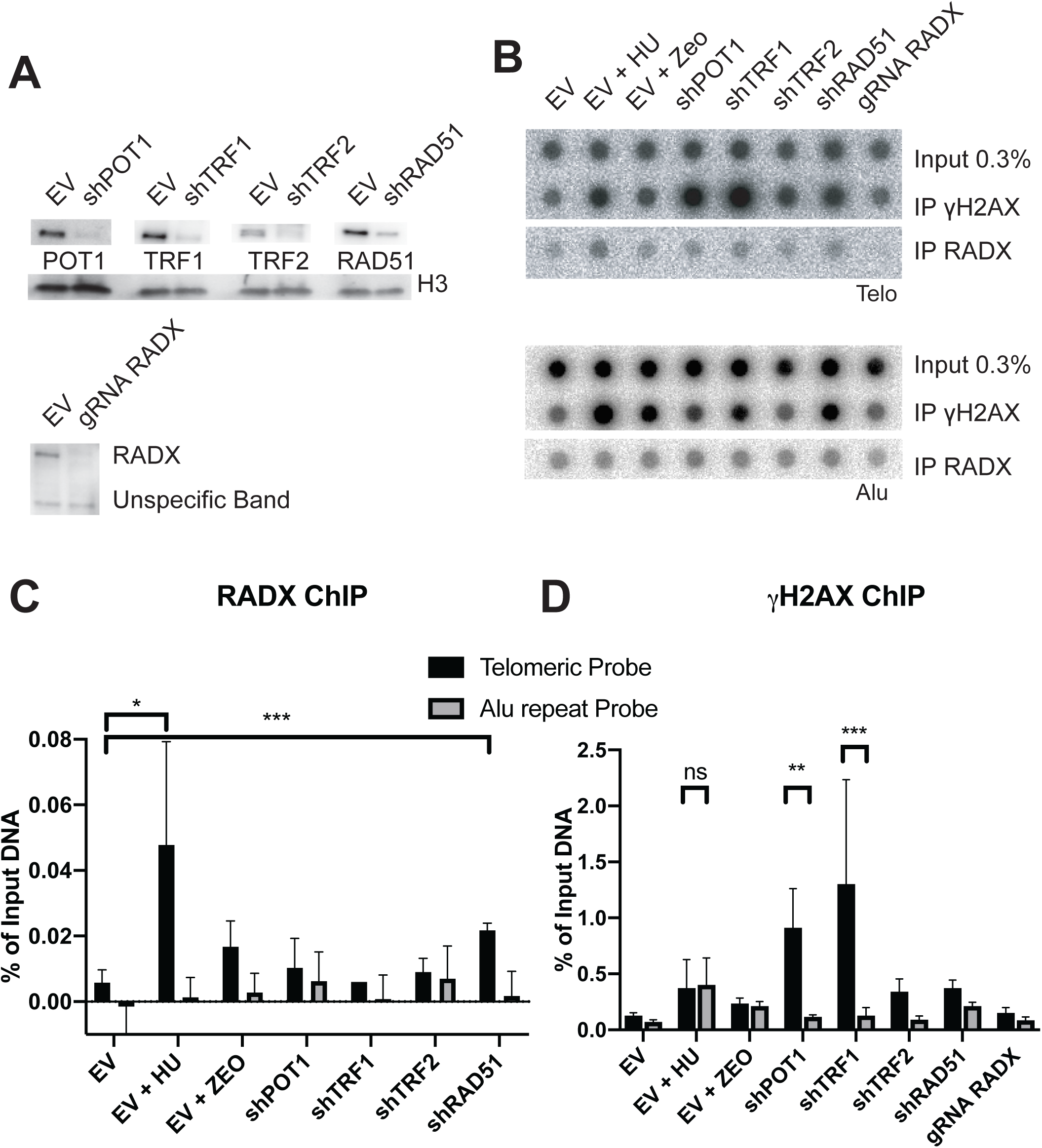
HU treatment increases RADX binding to telomeres. (A) Western blot of one representative experiment showing the efficiency of protein depletions in HEK293T cells. (B) Dot blot membranes of one representative experiment, hybridized with C-rich telomeric probe, stripped and rehybridized with Alu repeat probe. (C) and (D) Quantification of three independent ChIP experiments. Significance was determined by ordinary two-way ANOVA with multiple comparison test and p<0.05. % Input is displayed and in (C) the signal of the gRNA RADX sample was subtracted for every other sample to account for background binding. Cells were incubated with 1 mM HU or 1mM Zeocin for 14 h.

### RADX Cooperates with POT1 to Suppress RAD51-Dependent Telomere Fragility and Telomere Sister-Chromatid Associations

To uncover the biological functions of RADX at telomeres and its putative collaboration with the single-stranded telomeric DNA protein POT1 we carried out depletion studies and determined telomere integrity by staining telomeres in metaphase chromosomes by FISH. In a first series of experiments we depleted RADX with siRNAs in HEK293E cells containing conditional alleles of *POT1*, which could be deleted *via loxP*-sites upon expression of Cre-recombinase (Figure 4A; see accompanying paper by Glousker et al.). Telomere replication defects give rise to fragile telomeres (Sfeir et al., 2009). Telomeres were scored as fragile (white arrows in Figure 4C), when the telomeric signals from one chromosome arm were split into two or when the telomeric signal was elongated and diffuse. RADX depletion on its own did not change the levels of telomere fragility or sister-chromatid associations (Figures 4D and 4E; see red arrows in Figure 4C for sister-chromatid associations). *POT1* deletion increased telomere fragility as well as sister-chromatid associations consistent with published results (Pinzaru et al., 2016). Very strikingly, telomere fragility was drastically aggravated when RADX was depleted in *POT1*-knockout (KO) cells (Figures 4C and 4D). Telomere sister-chromatid associations also increased upon depletion of RADX in *POT1-KO* cells (Figures 4C and 4E). Co-depletion of TRF1 and RADX did not have the same effect (Figure S5). These results suggested that RADX and POT1 cooperate to suppress both of these telomere abnormalities. To test if the putative RADX antagonist RAD51 mediated these phenotypes, we co-depleted RAD51 or the RAD51 loader BRCA2. Indeed, the elevated telomere fragility as well as telomere sister chromatid associations were suppressed upon co-depletion of RAD51 or BRCA2 (Figures 4D and 4E). These results suggested that POT1 and RADX suppress homologous recombination at telomeres which when unleashed mediates sister-telomere associations and telomere fragility.

**Figure 4.**
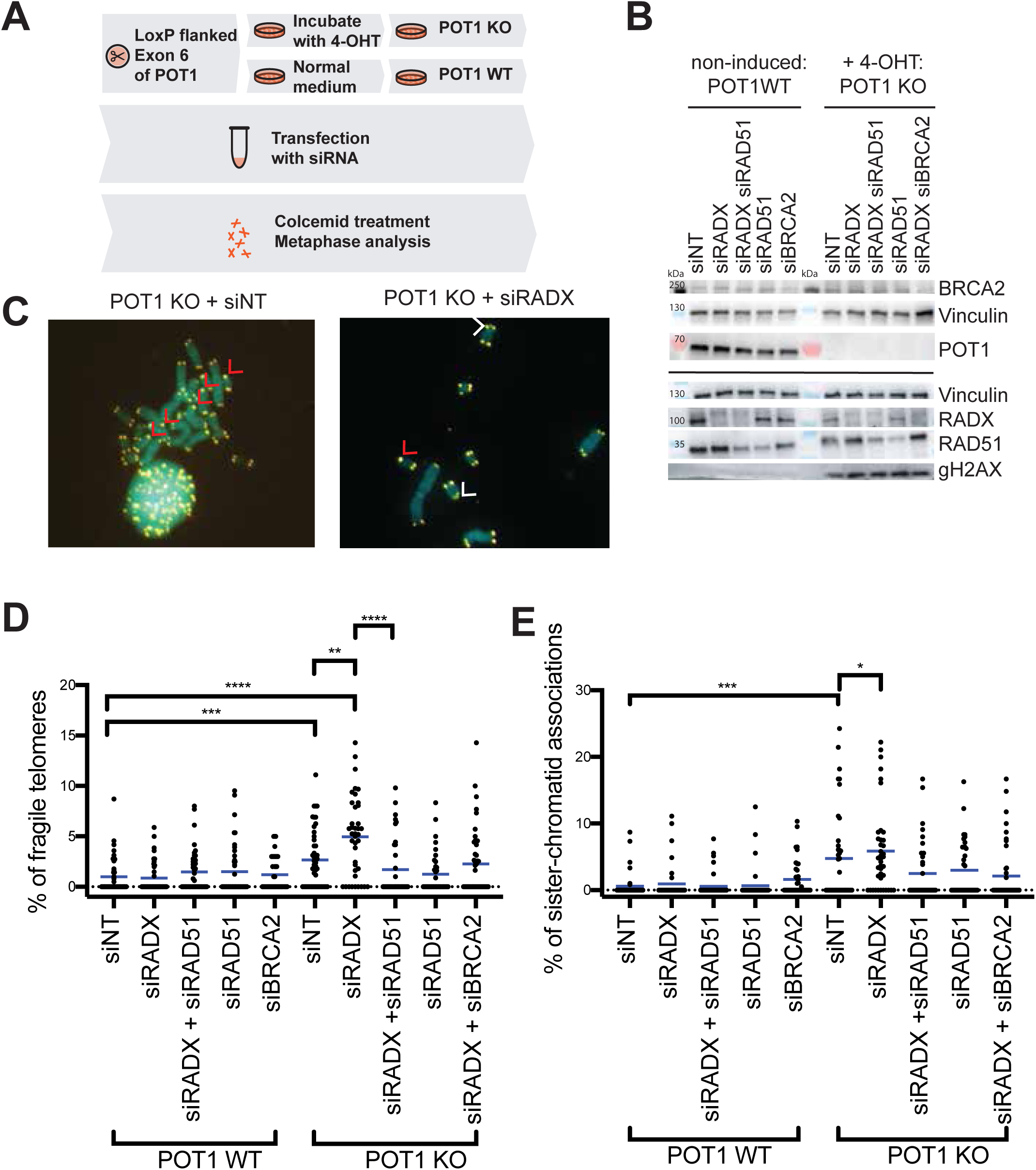
POT1/RADX- double depletion induces telomere fragility and telomere sister-chromatid associations. (A) Schematic explaining the experimental set-up for protein depletions in HEK293T cells. (B) Western blot of one representative experiment proving efficient gene deletions and protein depletions. SiRNAs were pools from Dharmacon. (C) Representative metaphases for induced POT1 KO cells transfected with non-targeting (NT) siRNA or siRADX. White arrows indicate fragile telomeres (smeary dot or two dots on one chromosome arm) and red arrows represent sister chromatid fusions. (D) and (E) Quantification of >37 metaphases per condition from three independent experiments. The mean is displayed and statistical significance was determined by ordinary one-way ANOVA and TUCKEY’S multiple comparison test, **** p<0.0001, *** p<0.005, ** p< 0.001 and * p<0.05.

To further corroborate these findings, we inverted the experimental design by first generating *RADX*-*KO* clones in HEK2913T cells using CRISPR/Cas9 and depleting POT1 with shRNAs (Figure 5A). Also, in this setting, telomere fragility was more pronounced upon depletion of POT1 in *RADX-KO* cells than in wild type cells and as above, telomere fragility was suppressed upon RAD51 co-depletion (Figures 5B, 5C and 5D). However, depletion of POT1 with shRNA in wild type and *RADX-KO* cells did not lead to an increase in sister-chromatid associations (Figure 5E). This therefore indicates that the small amounts of POT1, which were retained upon expression of shRNAs (Figure 5B) were sufficient to suppress telomere sister-chromatid associations. Still, these results confirmed that RADX and POT1 cooperate to suppress RAD51-dependent telomere fragility.

**Figure 5.**
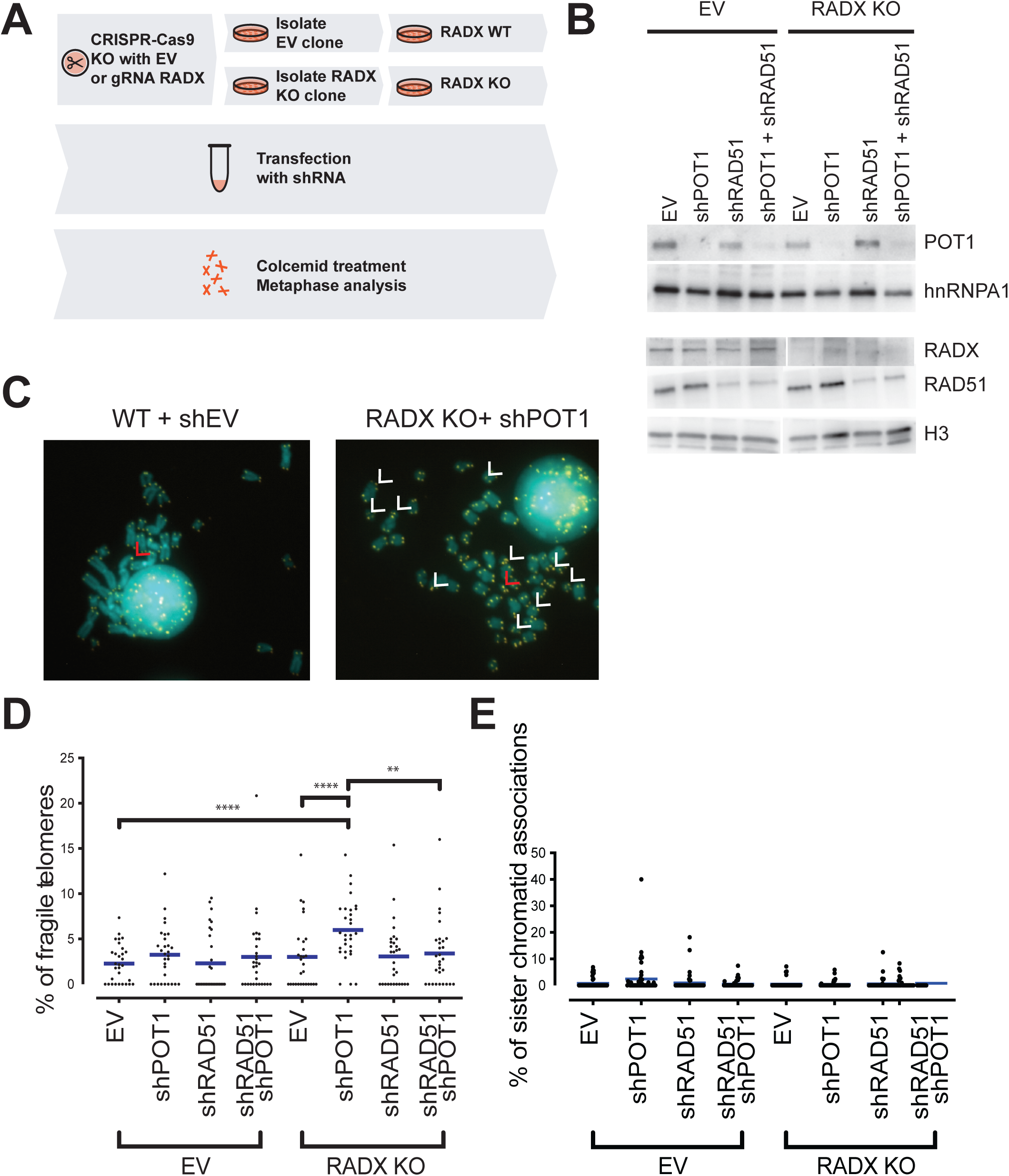
RADX/POT1- double depletions induce telomere fragility. (A) Schematic explaining the experimental set-up for protein depletions in HEK293T cells. (B) Western Blot of one representative experiment indicating efficient gene deletions and protein depletions. (C) Representative metaphases for induced POT1 KO cells transfected with non-targeting (NT) siRNA or siRADX. White arrows represent fragile telomeres (either a smeary dot or two dots on one chromosome arm) and red arrows represent sister chromatid fusions. (D) and (E) Quantification of >40 metaphases per condition from three independent experiments. The mean is displayed and statistical significance was determined by ordinary one-way ANOVA and TUCKEY’s multiple comparison test, **** p<0.0001, *** p<0.005, ** p< 0.001 and * p<0.05.

### RADX and POT1 Suppress RAD51 Binding

The above results demonstrated that RADX and POT1 suppress RAD51 dependent telomere damage. To elucidate the underlying mechanism, we tested the hypothesis that RADX and POT1 suppress RAD51 binding at telomeres. We depleted POT1 via shRNAs and deleted *RADX* with CRISPR/Cas9 in Hela cells (Figure 6A). RAD51 association with telomeres was determined by quantifying colocalization of RAD51 with telomeres in IF/FISH experiments (Figure 6B). Deletion of *RADX* on its own did not significantly increase RAD51 abundance at telomeres (Figure 6B and 6C). However, POT1 depletion enhanced RAD51 binding which was further enhanced upon POT1 depletion in *RADX-KO* cells. This experiment therefore indicates that RADX and POT1 cooperate to suppress telomere association of RAD51.

**Figure 6.**
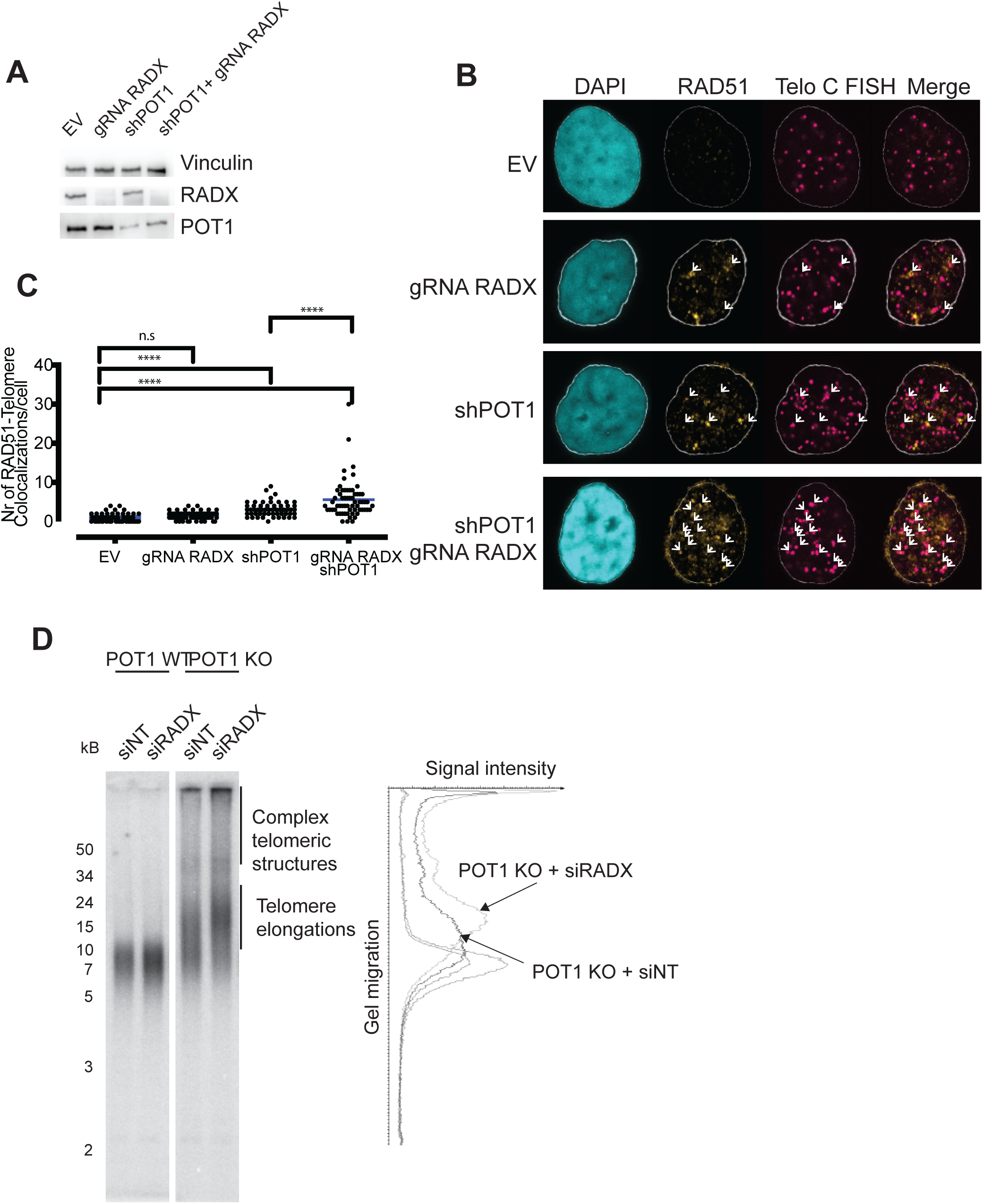
RADX and POT1 suppress RAD51 binding. (A) Western blot of one representative experiment showing efficient protein depletions in Hela cells. (B) Representative IF-FISH images of RADX or/and POT1 depletions with RAD51 in yellow and telomeres in pink. White arrows indicate colocalization. (C) Quantification of one representative experiment with >55 cells analyzed per condition. The mean is displayed and statistical significance was determined by ordinary one-way ANOVA and DUNNETT’s multiple comparison test, **** p<0.0001 and * p<0.05. (D) TRF of HEK293T cells with the same experimental set-up as for Figure 4, except for the skipping of demecolcine treatment.

Finally, we determined if *RADX*-deletion affected the telomeric 3’ overhang structure or telomere length homeostasis. The telomeric 3’ overhang was detected by Southern hybridization of non-denatured DNA and telomere length was determined by Southern hybridization of denatured DNA (Figure S6). In both of these analyses, *RADX* deletion had no impact on telomeric DNA length and structure. However, in the accompanying paper (Glousker et al.) we discovered the *POT1* deletion leads to rapid telomere elongation within seven days of growth. We therefore tested if RADX depletion would affect this phenotype and saw enhanced telomere elongation (Figure 6D) in cells deprived of POT1 and RADX indicating the cooperation of RADX and POT1 in also suppressing this phenotype.

### RADX is Important to Sustain ATR-signaling upon POT1-loss

Since RADX suppressed RAD51-binding to telomeres upon POT1 loss, we determined the effects of RADX/POT1-codepletion on ATR-dependent DNA checkpoint-signaling. We compared on Western blots the levels of the ATR substrates p-RPA32 (S4/8) and p-CHK1 (S345) in *POT1* wild type and *POT1*-KO cells upon co-depletion of RADX and RAD51 (Figure 7A). Consistent with previous data (Denchi and de Lange, 2007), we observed upon *POT1* deletion increased ATR-signaling shown by higher p-RPA- and p-CHK1-levels. The checkpoint response was significantly lowered if RADX was co-depleted. Concomitant depletion of RADX with RAD51 gave a similar checkpoint response as wild type cells whereas RAD51 depletion increased the checkpoint response significantly when compared to RADX depleted cells (Figure 7A). These results suggest that ATR-checkpoint-signaling is compromised when RADX is lost in addition to POT1, likely due to the BRCA2-mediated exchange of RPA against RAD51. On the other hand, RADX depletion did not perturb ATR-signaling upon global replication damage induced by HU treatment (Figure S7).

**Figure 7.**
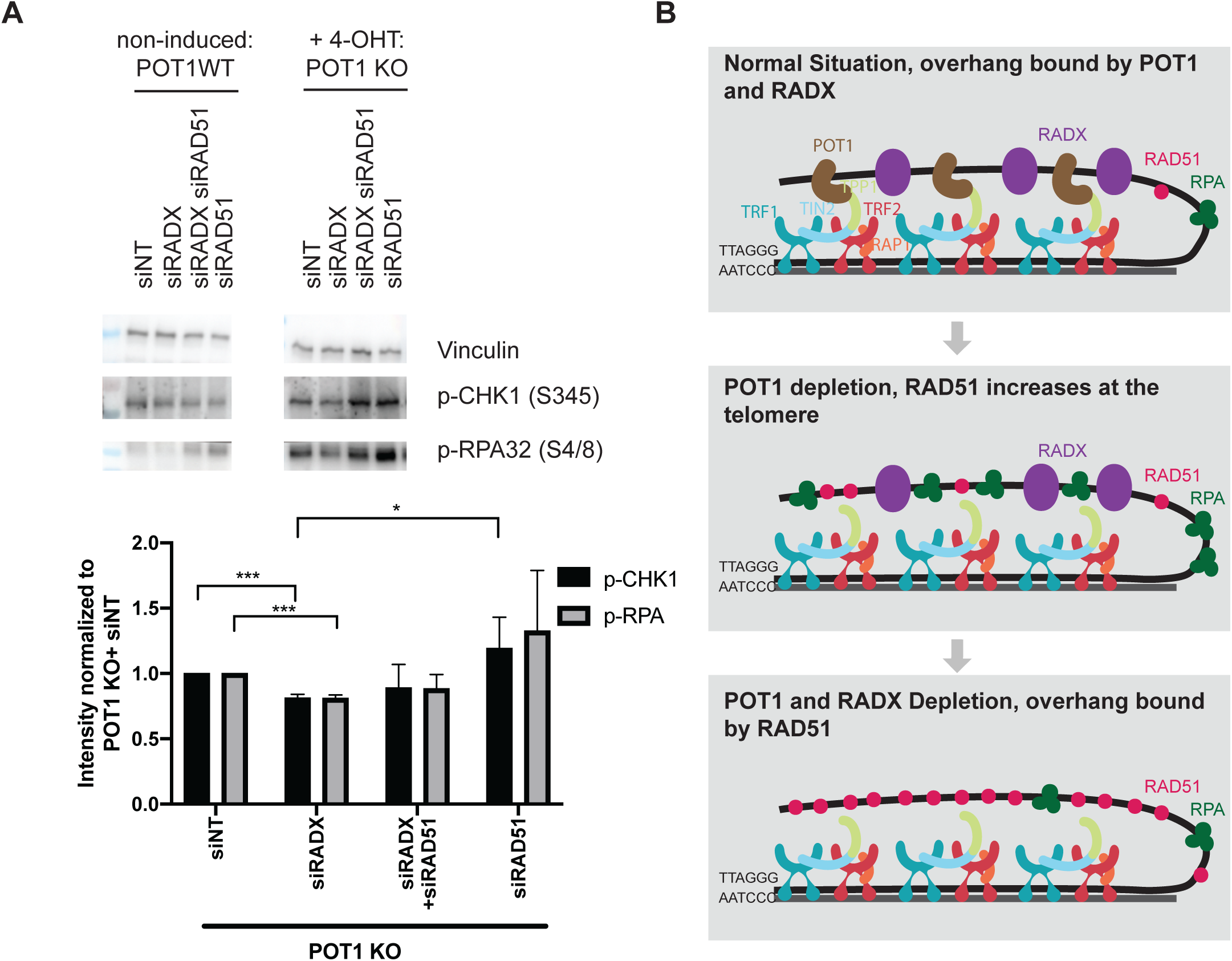
RADX is important to sustain ATR signaling upon POT1 loss. (A) Western blot of one representative experiment monitoring ATR-checkpoint signaling upon POT1 loss by blotting for p-ChK1 (S345) and pRPA (S4/8). Vinculin serves as loading control. Efficient protein depletions are shown in Figure 4B. Three independent experiments were quantified and p-CHK1 and pRPA levels were normalized to POT1+ siNT. Statistical significance was calculated by STUDENTS-t-Test, *** p<0.001. (B) Schematic model of RADX, POT1 and RAD51 binding at the telomeric overhang.

## Discussion

In this paper we explore the telomeric protein environment of TRF1, TRF2 and POT1 by inserting the BirA sequence at the endogenous loci in frame with the ATG start codons. The fusions did not interfere with telomere function when assessing telomere length maintenance and telomere integrity visualized on metaphase chromosomes. The overlap of the biotinylated proteins identified with all three shelterins was remarkable providing confidence in the biological relevance of identified proteins. Our live-cell labeling results are also consistent with previous data showing a physical association of shelterin components and the formation of functional complexes (Houghtaling et al., 2004; Liu et al., 2004; Ye et al., 2004). Still, slightly more proteins were identified with TRF1 and especially with TRF2 than with POT1 suggesting that the double strand telomere binding proteins may connect to a larger set of cellular processes.

### RADX Association with Telomeres

During the course of our studies CxORF57 was identified as an antagonist of RAD51 and re-named RADX (Dungrawala et al., 2017; Schubert et al., 2017). We focused our attention on RADX as it was identified with all three fusion proteins and because it contained OB-folds which are present not only in RPA but also in the important telomere components POT1, TPP1 and CST complex subunits. Our data indicate that RADX binds along with POT1 to the single-stranded G-rich strand at telomeres which is present as 3’ overhang or present as displaced strand when telomeres are engaged in t-loops. RADX binding occurs in S-phase as well as interphase. This conclusion is supported by the observation that telomere association is dependent on an intact single-strand-DNA-binding-domain present in OB2 and independent of S-phase. Furthermore, telomere binding was enhanced upon HU treatment, which results in replication fork stalling. In this experiment we observed DNA damage signaling at telomeres and at Alu repeats, but RADX was enriched only at telomeric repeats suggesting there may be preferential binding of RADX to telomeric sequences. Interestingly, TRF1 depletion, which previously was also shown to increase replication fork stalling and replication problems at telomeres (Sfeir et al., 2009) did not induce a comparable rise in RADX telomere binding. We hypothesize that RADX binds specifically to unmanteled G-rich DNA, which is free upon replication fork stalling but forms G4 structures upon TRF1 removal mediated replication fork stalling due to lack of helicases like BLM which resolve G4 structures (Zimmermann et al., 2014).

### RADX Cooperation with POT1 to Suppress Telomere Fragility and Sister-Chromatid Associations

Our results demonstrate that co-depletion of RADX with POT1 but not with TRF1 enhances telomere fragility and sister-telomere associations. It was recently demonstrated that RADX counteracts RAD51 mediated replication fork reversal (Dungrawala et al., 2017) which occurs independently of BRCA2 (Mijic et al., 2017). We tested the involvement of RAD51 and BRCA2 for POT1/RADX depletion mediated telomere fragility and sister-chromatid associations and found that both, RAD51 or BRCA2 depletion, rescued the telomere damage. This suggests that telomere fragility and sister-telomere associations are generated due to activation of homologous recombination, rather than fork reversal at telomeres.

Altogether, our results support a model in which RADX represents next to the more abundant POT1 a second OB-fold containing telomere binding protein that contributes to telomere protection (Figure 7B). Concomitant loss of POT1 and RADX leads to efficient BRCA2-mediated loading of RAD51 at telomeres unleashing RAD51 mediated sister-chromatid association and HDR resulting in telomere fragility. When POT1 is abundantly present at telomeres, the telomeric RADX function is not yet apparent. POT1 is capable to suppress RAD51 binding and the resulting telomere abnormalities on its own, independently of RADX. However, when POT1 becomes scarce at telomeres, RADX function becomes critical to support POT1 in suppressing RAD51 binding (Figure 6B). This function is important also to sustain RPA-ATR-ATRIP mediated damage signaling at telomeres preventing the exchange of RPA by RAD51 (Figure 7A). We predict that this mechanism becomes particularly critical during cellular senescence when POT1 concentration is lowered at telomeres to guarantee long-lasting RPA-ATR/ATRIP mediated checkpoint signaling. Thus, our results should inspire future investigations to test the roles of RADX for the maintenance of cellular senescence to suppress tumorigenesis.

## Author contributions

A-SB carried out all experiments except for the development of the *POT1* conditional knockout cell line, which was done by GG. A-SB and JL wrote the paper. All authors conceptualized the experiments.

## Acknowledgements

We thank Timothy Murray for collecting samples and performing the TRF analysis in Figure S6, as well as Chih-Yi Lin for help with proteomic comparison analysis in Figure S3B. We further thank Romain Hamelin and the proteomics core facility at EPFL for mass spectrometry analysis. We are grateful to Thomas Lunardi for technical help. Current and former members of the Lingner lab are thanked for discussions, technical advice and sharing reagents. Research in J.L.’s laboratory was supported by the Swiss National Science Foundation (SNSF), the SNSF funded NCCR RNA and disease network, an Initial Training Network (ITN) grant (aDDRess) from the European Commission’s Seventh Framework Programme, the Swiss Cancer League and EPFL.

## Supplemental Information

Supplemental Information includes figures S1-S6 as well as table S1.

**Figure S1.**
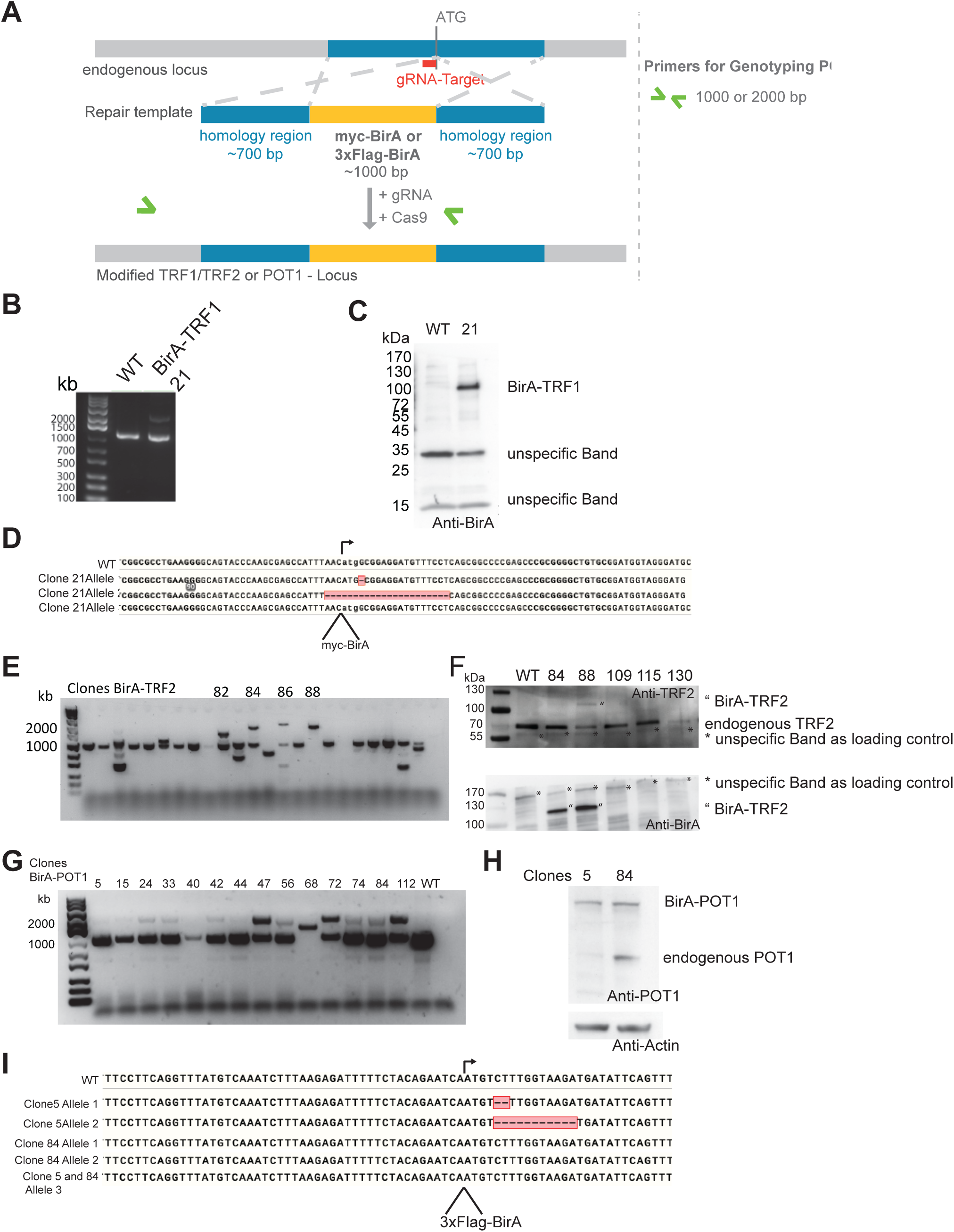
Screening and characterization of BirA-clones. (A) Schematic explaining the PCR approach for screening and genotyping of BirA-tagged clones. (B) Amplification products for the genotyping PCR showing a band for an untagged WT allele and a larger band for the integration of the BirA sequence at the endogenous TRF1 locus in clone 21. (C) Western blot showing the expression of BirA-tagged TRF1 in clone 21. (D) Sequencing results of the subcloned PCR products from (C) for clone 21. (E) Amplification products for the genotyping PCR for BirA-TRF2 clones. (F) Western blots for selected BirA-TRF2 clones and a WT control. (G) Genotyping PCR for BirA-POT1 clones. (H) Western blot for BirA-POT1 clone 5 and 84 showing endogenous and tagged POT1. (I) Sequencing results for BirA-POT1 clones 5 and 84.

**Figure S2.**
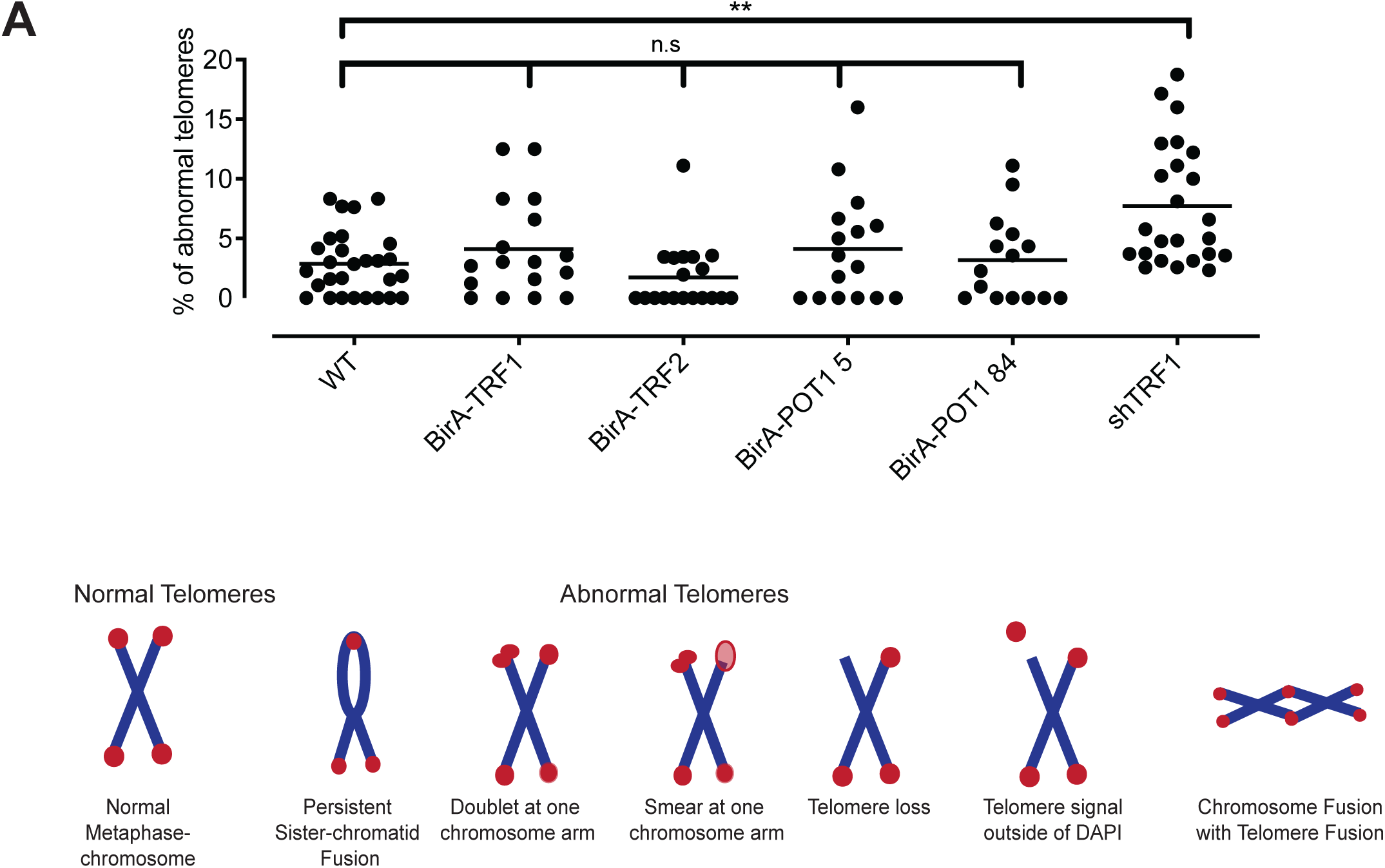
Functional characterization of BirA-clones. (A) WT and BirA clones derived metaphase chromosomes were analyzed for abnormal telomeres and compared to TRF1 depletion as positive control. Statistical significance was determined by one-way-ANOVA and DUNETT’s multiple comparison test and **p<0.01%.

**Figure S3.**
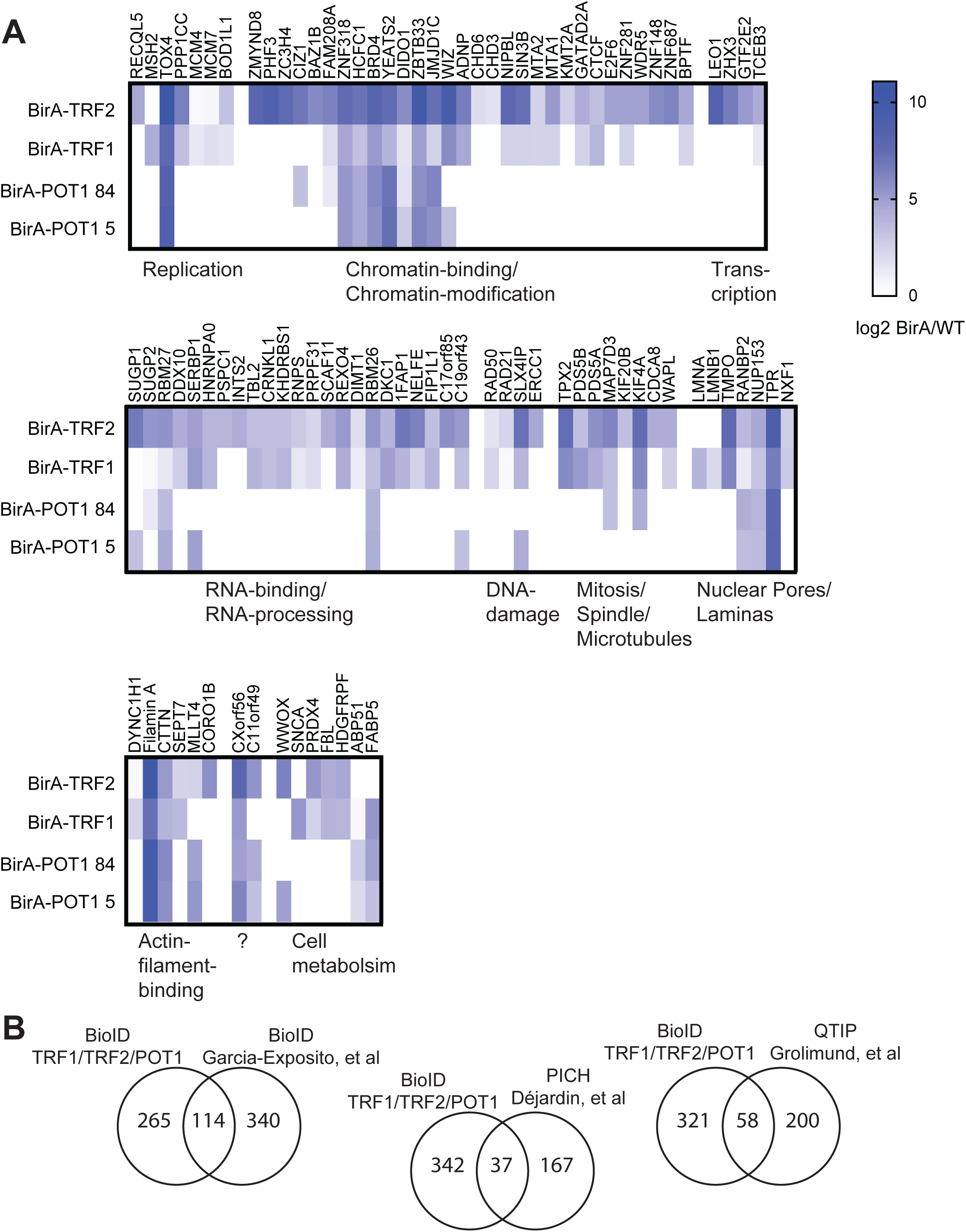
Identified proteins in BioID. (A) Heatmap showing enrichment ratios, calculated by log2 ratios of total spectral counts BirA/WT, for a selection of proteins identified in the corresponding BioIDs. Proteins were not identified in the WT control IP or at least 3-fold enriched (total spectrum counts) in BirA-IP and identified in at least two BirA-IPs. (B) Comparison of proteins identified in this study to telomeric proteins identified by PICH (Déjardin and Kingston, 2009), TRF1-BioID (Garcia-Exposito et al., 2016) or QTIP (Grolimund et al., 2013).

**Figure S4:**
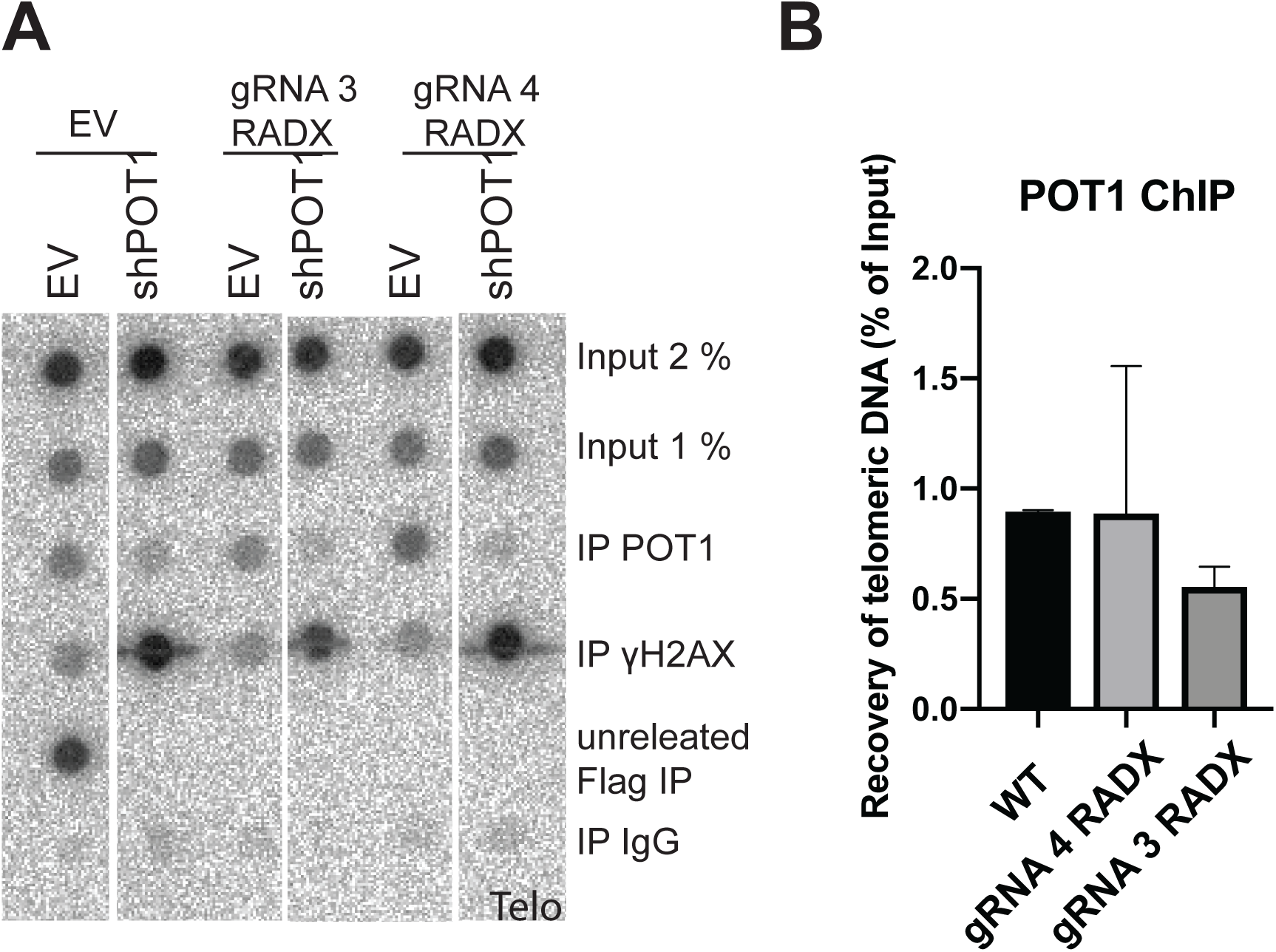
RADX depletion does not change POT1 abundance at the telomere. (A) Dot blot membranes of one representative experiment carried out with HEK293T cells and hybridized with a C-rich telomeric probe. (B) Quantification of three independent ChIP experiments. Significance was determined by ordinary one-way ANOVA with DUNNETT’s multiple comparison test and p>0.05.

**Figure S5.**
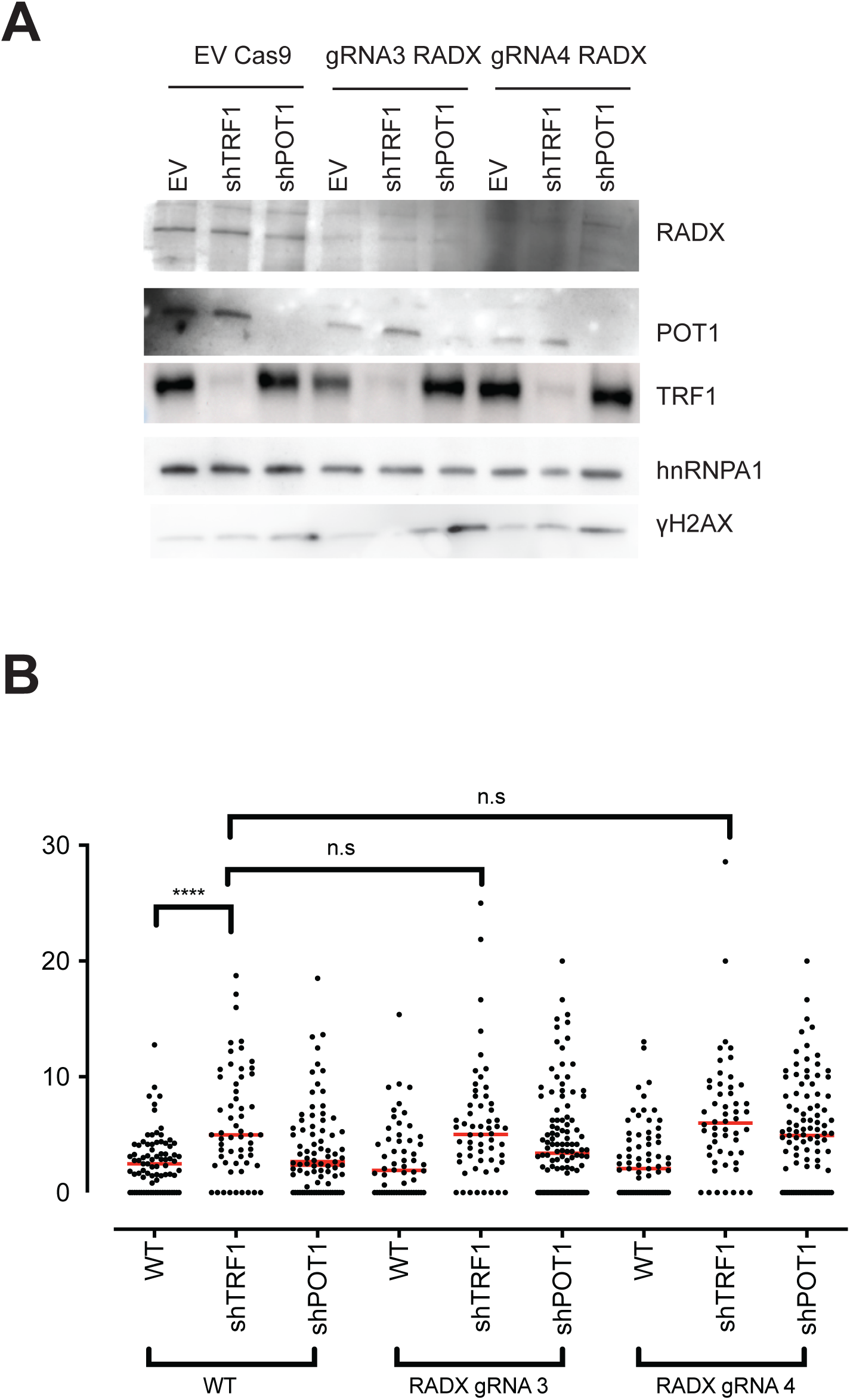
RADX/TRF1 double depletion does not increase telomere fragility. (A) Western blot of one representative experiment indicating efficient gene deletions and protein depletions in Hek293T cells. (B) Quantification of >90 metaphases per condition from three independent experiments. The mean is displayed and statistical significance was determined by ordinary one-way ANOVA and TUCKEY’s multiple comparison test, **** p<0.0001, *** p<0.005, ** p< 0.001 and * p<0.05.

**Figure S6.**
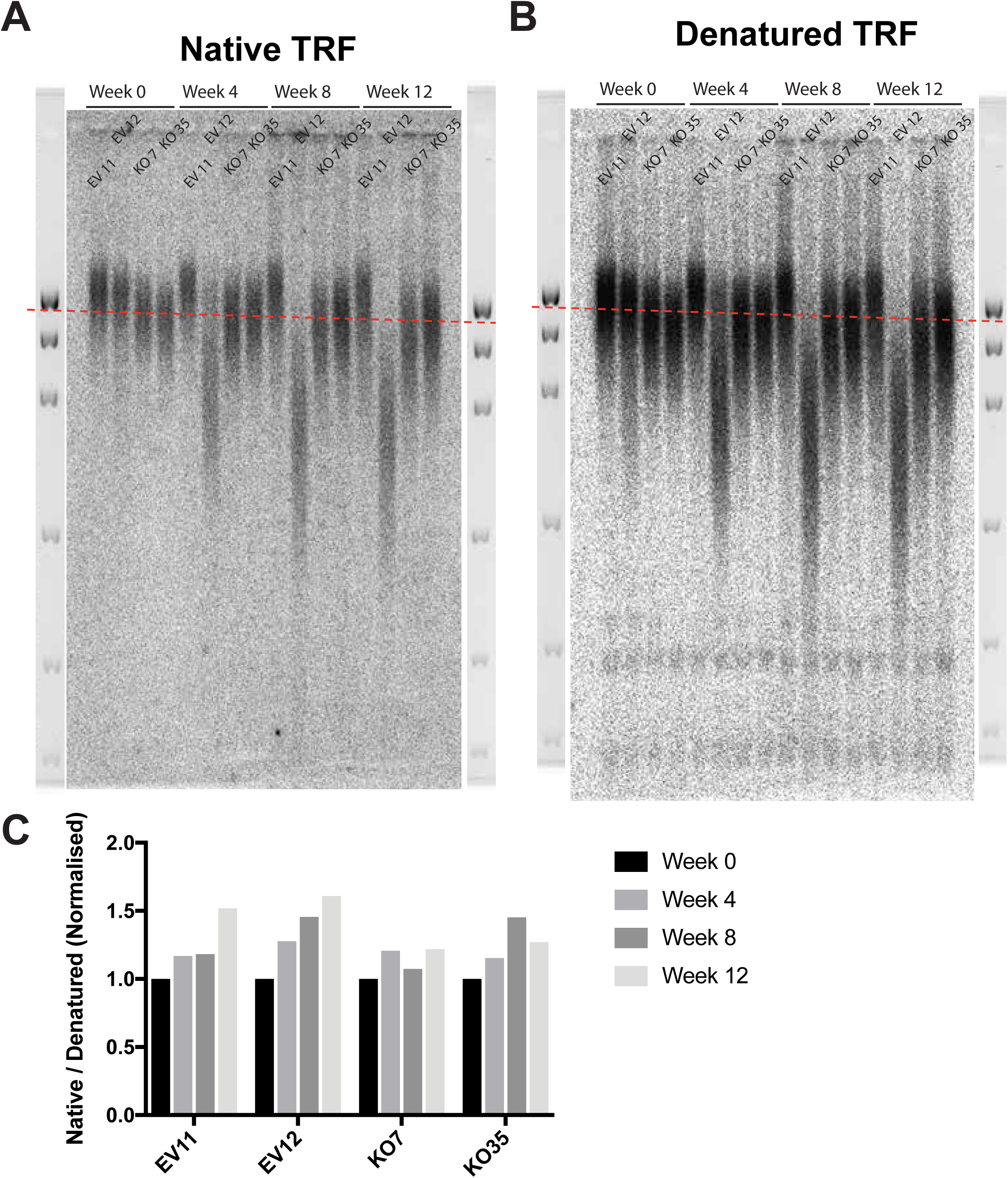
RADX KO does not change telomeric 3’overhang and total telomere length. (A) + (B) Telomere restriction fragment analysis of two isolated HEK293T clones transfected with EV, gRNA3 RADX (KO7) or gRNA4 (KO35). The same gel was hybridized with a 32P-labelled Telo-C-probe with or without denaturation. (C) Quantification of the native/denatured signal normalized to week 0 from the gel in (A) and (B).

**Figure S7.**
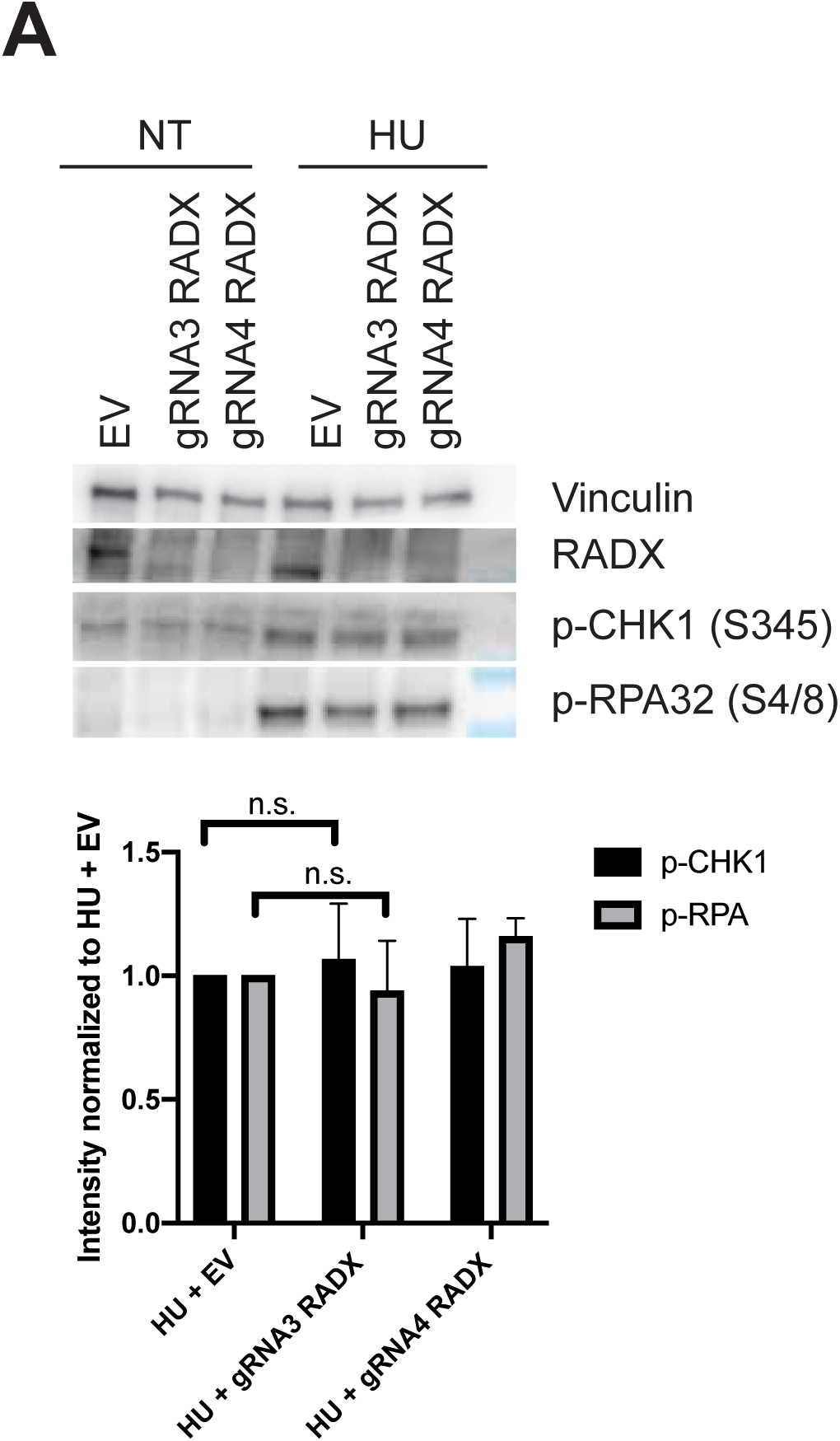
RADX is not important to sustain ATR-checkpoint signaling upon general replication stress. (A) Western blot of one representative experiment monitoring ATR-checkpoint signaling upon HU 1mM incubation for 14h by blotting for p-ChK1 (S345) and pRPA (S4/8). Vinculin serves as loading control. Three independent experiments were quantified and p-CHK1 and pRPA levels were normalized to EV + HU. Statistical significance was calculated by STUDENTS-t-Test, n.s. p>0.05.

## Star Methods

### Experimental Procedures

#### Cell culture

Hela and HEK293T cells were cultured in Dulbecco’s modified Eeagle’s medium supplemented with 10% fetal calf serum and 100 U/ml of penicillin/streptomycin. Cells were grown at 37°C in presence of 5% CO_2_.

#### Lipofectamine transfection

0.7 Million HEK293T cells were seeded the day before transfection in a 6-well plate in 2 ml DMEM/FBS/PenStrep. Cells were transfected with 3 μg DNA per plasmid, suspended in 250 μl OptiMEM, combined with 10 μl Lipofectamie 2000 (Invitrogen) in 250 μl OptiMEM. After 15 min incubation the transfection mix was added dropwise to the cells. After 18-24 h the cells were split and transferred to a 10 cm dish and incubated with either blasticidin (5 μg/ml) or puromycin (1 μg/ml) for 3 days. Cells were split again, fresh medium DMEM/FBS/PenStrep was added and cells were harvested after 1-3 days.

#### siRNA transfection in POT1-inducible KO cells

HEK293E cells were induced with 0.5 μM 4-Hydroxytamoxifen on day 0. On day 2, cells were split and 5 million cells were seeded into 10 cm dishes. On day 3 the medium was changed to 5 ml DMEM/FBS without PenStrep and transfected with 30 pmol siRNA (222 μl H_2_O, 25 μl 2.5 M CaCl_2,_ 250 μl 2x HBSS pH 7.04, 3 μl 10 μM siRNA). On day 4, the medium was changed and cells were split into 2-3 10 cm dishes. Cells were harvested on day 6.

#### Plasmid preparation

For validation of telomeric localization of RADX by IF-FISH, RADX cDNA was amplified from HEK293T cells and cloned into pcDNA6 downstream of three tandemly repeated Flag tags giving rise to pcDNA6-3xFlagRADX. The shRNA expressing vectors were created by ligating the double-stranded DNA oligonucleotides into pSuper-puro or pSuper-Blast-vectors (Oligoengine) digested with BglII and HindIII. gRNAs were generated by ligating the annealed DNA oligonucleotides into the BbsI restriction site of pSpCas9(BB)-2A-Puro (Addgene 48139) as described (Ran et al., 2013). RADX *OB was generated by site-directed-mutagenesis of pcDNA6-3xFlagRADX (QuickChange II Site-directed mutagenesis kit, Agilent).

#### CRISPR/Cas9 gene editing BirA clone generation

gRNAs to target the genomic loci in proximity to the start codon were selected with the tool provided by Feng Zheng’s laboratory (Broad institute, (Shalem et al., 2014)) and were cloned into pSpCas9 (BB)-2A-GFP as described (Ran et al., 2013). The repair template was created by amplifying the homology regions from gDNA of HEK293T cells and the BirA sequence from pcDNA3.1-myc-BioID (Addgene 35700) and combined by overlapping PCR (Bryksin and Matsumura, 2010) and further subcloned into TOPO Zero Blunt for Sequencing (Invitrogen). 1 Million Hek293T cells were transfected in a 6-well plate containing 2 ml DMEM/FBS with 3 μg DNA gRNA and 3 μg repair template, 500 μl OptiMEM and 10 μl Lipofectamine. Cells were split the next day and after 72 h single GFP-positive cells were sorted into 96-well plates. After 2 weeks, cells were split into two 12-well plates. Three days later genomic DNA was extracted from one row of 12 well plates with the Wizard genomic DNA Purification System (Promega). PCR with primers flanking the region of insertion was performed to screen for clones with insertion of the BirA sequence. Positive clones were further expanded and the PCR product was subcloned into TOPO Zero Blunt for sequencing and genotyping.

#### RADX KO clone generation

gRNAs were selected based on low off-target scores with the tool from Feng Zheng’s laboratory (Broad institute, (Shalem et al., 2014)) and cloned into pSpCas9 (BB)-2A-Puro as described before. 1 Million Hek293T cells were transfected in a 6-well plate containing 2 ml DMEM/FBS with 4 μg DNA, either gRNA or EV plasmid, 500 μl OptiMEM and 10 μl Lipofectamine. The next day cells were split and puromycin (1 μg/ml) was added. After 3 additional days cells were diluted to single cells and 1 cell per well was seeded in a 96 well plate. After 2.5 weeks colonies were screened for RADX loss by western blot and validated by sequencing.

#### Western blot

0.5 million cells were incubated in 2x Laemmli buffer with 100 mM DTT and boiled at 95°C. Proteins were separated on a 4-15% gradient Mini Protean TGC (BioRAD) followed by wet transfer onto a 0.2 μm nitrocellulose membrane (Amersham Protran, GE Healthcare). Membranes were blocked either in 3% BSA in 1x PBS+ 0.1% Tween or 5 % milk in 1x PBS + 0.1% Tween overnight and washed 3x 15 min with PBS + 0.1% Tween the next day. HRP-conjugated secondary antibodies in combination with ECL spray (Advansta) were used to reveal the signal on a FluorChem 8900 (Alpha Innotech) or Fusion FX (Vilber) detector.

#### Telomere restriction fragment analysis (TRF)

Genomic DNA from 3 million cells was isolated with the Wizard genomic DNA kit (Promega) following the protocol of the manufacturer. 5 μg genomic DNA was digested with 15 U RsaI and 15 U HinfI (New England Biolabs) overnight at 37°C. Samples were loaded on a 15 cm long 0.8% agarose gel and separated. Alternatively, DNA was separated by pulse field gel electrophoresis on a 1% agarose gel in 0.5 x TBE at 5 V cm^-1^ for 16 h at 14 °C with switch times ramped form 0.5 to 6 seconds. After electrophoresis, the gels were dried for 2h at 50°C, denatured with 0.8 M NaOH, 0.5 M NaCl, neutralized with 0.5 M Tris-HCl pH 7.0 and prehybridized at 50°C in Church buffer (1% BSA, 7% SDS, 1 mM EDTA, 0.5 M Na-phosphate buffer at pH 7.2). Hybridization was overnight at 50°C with a ^32^P-labeled telomeric probe as described (Grolimund et al., 2013). After hybridization the gel was washed for 1 h with each wash buffer at 50°C containing 2x SSC + 0.1% SDS, 1x SSC + 0.1% SDS, 0.5x SSC + 0.1% SDS and 0.1x SSC. Radioactive signal was detected with Amersham Typhoon.

#### Telomeric FISH on metaphase chromosomes

Slides were prepared as described (Majerska et al., 2018).

#### ChIP-dot blot

15 million HEK293T cells were crosslinked for 15 min at room temperature in 1.5 ml 1 % methanol-free formaldehyde in PBS. Formaldehyde was quenched for 5 min with 0.1 M glycine for 5 min. Chromatin enriched fractions were prepared by lysis in SDS lysis buffer (1% SDS, 50 mM Tris-HCl pH 8.0, 10 mM EDTA pH 8.0). Chromatin was resuspended in 1 ml LB3 (10 mM Tris-HCl pH 8.0, 200 mM NaCl, 1 mM EDTA, 0.5 mM EGTA, 0.1 % Na-deoxycholate, 0.25% sodium lauroyl sarcosinate, EDTA-free protease inhibitor complex (Roche) and sonicated for 15 min at 4°C using a Focused-Ultrasonicator (Covaris, E220, duty 5.0, PIP: 140, cycles: 200, amplitude 0, velocity 0, dwell 0, 0.12 x12 mm glass tubes with AFA fiber). Insoluble material was removed by centrifugation for 15 min, 4°C, at 20,000 g. Sonicated extracts were diluted with 5 volumes IP dilution buffer (50 mM Tris-HCl pH 8.0, 0.75% Triton-X-100, 600 mM NaCl, 10 mM EDTA pH 8.0) and precleared with 30 ul sepharose protein G beads (GE Healthcare), which were blocked with 1 mg/ml BSA. Immunoprecipitation was performed with 30 μl blocked sepharose protein G beads and 5 μl of antibody or serum per precleared lysate corresponding to 200,000 cells. After overnight incubation at 4°C, beads were washed at 4°C for 5 min on a rotating wheel with the following washing buffers: Wash 1 (0.1% SDS, 1% Triton-X-100, 2 mM EDTA pH 8.0, 20 mM Tris-HCl pH 8.0, 300 mM NaCl) for 5 min, wash 2 (0.1% SDS, 1% Triton-X-100, 2 mM EDTA pH 8.0, 20 mM Tris-HCl pH 8.0, 500 mM NaCl) for 5 min, wash 3 (250 mM LiCl, 1% NP-40, 1% Na-deoxycholate, 1 mM EDTA pH 8.0, 10 mM Tris-HCl pH 8.0) for 5 min and twice with TE. Beads were suspended in 100 μl crosslink reversal buffer (20 mM Tris-HCl pH 8.0, 1% SDS, 100 mM NaHCO_3,_ 0.5 mM EDTA, 200 μg/ml RNase-DNase free (Roche)) and incubated 5 h or overnight at 65°C. DNA was purified using the NucleoSpin Gel and PCR Clean-up with buffer NTB (Macherey Nagel) and eluted in 100 μl TE buffer. Afterwards DNA was denatured 5 min at 95°C and chilled immediately on ice before spotting onto a Hybond N+ nylon membrane (GE Healthcare) using a BioRad dot blot apparatus. The membrane was UV-crosslinked, denatured with 0.8 M NaOH, 0.5 M NaCl and neutralized with 0.5 M Tris-HCl pH 7.0 and blocked in Church buffer at 65°C for 1 h. Incubation with a ^32^P-labeled telomeric probe was done overnight at 65 °C as described (Grolimund et al., 2013). The next day the membrane was washed 3x 30 min with 1x SSC + 0.5% SDS. For Alu probe ((5′-TGATCCGCCCGCCTCGGCCTCCCAAAGTG-3′) incubation, membranes were stripped by boiling at 95°C for 3x 10 min in 0.1x SSC + 1% SDS. Membranes were again prehybridized with Church buffer at 55 °C and hybridized with the Alu repeat probe in Church buffer overnight at 55°C and washed 3x 30 min with 1x SSC + 0.5% SDS.

Radioactive signal was detected with a FujiFilm Fluorescent Image Analyzer (FLA-3000) and the intensity of each dot was calculated using AIDA software. Averages and p-values were calculated using PRISM 8 software.

#### IF-FISH

Hela cells were grown on coverslips, eventually incubated with 10 mM EdU in DMEM for 10 min at 37°C, washed with PBS, incubated for 7 min on ice with pre-extraction buffer (0.5% Triton-X-100, 20 mM HEPES/KOH pH 7.9, 50 mM NaCl, 3 mM MgCl_2_, 300 mM sucrose) and fixed with 4% formaldehyde. Then cells were permeabilized for 5 min in 0.1% Triton-X-100/0.02 % SDS/ 1x PBS and pre-blocked with 2% BSA in PBS for 10 min before blocking with 10% goat-serum/ 2 % BSA /1x PBS for 45 min. For EdU-analysis, cells were labeled with a Click-it reaction (20 mM Copper sulfate, 100 mM sodium ascorbate, 2 mM Alexa 488-azide (Invitrogen), PBS 1x) for 30 min before blocking. Afterwards cells were incubated with primary antibody in blocking solution for 90 min, washed 3x with 2% BSA/PBS and stained with secondary antibody in 2% BSA/PBS for 45 min. Further, cells were washed 3x times with PBS and fixed in 4% formaldehyde for 5 min. For telomeric FISH, coverslips were dehydrated in ethanol series, air dried and hybridized with a Cy3-OO-(CCCTAA)_3_ PNA probe as described (Majerska et al., 2018). Images were taken with a Zeiss LSM 700 confocal microscope equipped with a 63x oil immersion objective.

#### BioID

800 million HEK293T cells expressing BirA-TRF1/TRF2 or POT1 were grown in normal Dulbecco’s modified Eeagle’s medium supplemented with 10% fetal calf serum and 100 U/ml of penicillin/streptomycin. Biotin (Sigma) was added to a final concentration of 50 μM for 24 h. Cells were harvested with trypsin and washed with 1x PBS. Cells were resuspended in 40 mL buffer A (20 mM HEPES, pH 7.5, 10 mM KCl, 1.5 mM MgCl_2_, 0.34 M sucrose, 10 % glycerol, 1 mM dithiothreitol, 0.2% Triton-X-100, and protease inhibitors (Roche)) and incubated on ice for 5 min. Nuclei were pelleted by centrifugation at 600 g for 5 minutes at 4°C. Then nuclei were washed three times with 200 mL buffer M (10 mM HEPES, pH 7.5, 60 mM KCl, 10 mM NaCl). Pellets were resuspended in 30 mL RIPA buffer (0.5% Na-deoxycholate, 0.1% SDS, 1% NP-40, 150 mM NaCl, 1 mM EDTA, 1mM EGTA, 50 mM Tris-HCl) supplemented with protease inhibitors (Roche) and 150 U/mL of benzonase (Sigma). Samples were incubated for 1 h at 4°C on a rotating wheel. The insoluble fraction was separated by centrifugation at 18,500 g for 15 min at 4°C. The protein concentration of the supernatant was determined using the Bradford assay (Bio-Rad). The amount of protein was adjusted between the different samples. 100 mg of protein and 660 μl of magnetic streptavidin beads (Dynabeads MyOne Streptavidin C1, Thermo Fischer Scientific) were used for the IP. Beads were washed three times with RIPA buffer before being added to the nuclear extracts. Samples were incubated on a wheel overnight at 4°C. Beads were washed for 5 minutes with wash 1 (2% SDS), wash 2 (0.1% Na-deoxycholate, 1% Triton, 1 mM EDTA, 500 mM NaCl, 50 mM HEPES), and twice with wash 3 (0.5 % Na-deoxycholate, 0.5% NP-40, 1 mM EDTA, 250 mM LiCl, 10 mM Tris-HCl pH 8.0). Beads were resuspended in 50 μL of 4x Laemmli buffer with 355 mM of β-mercaptoethanol and run on a 10% Mini Protean TGC (BioRAD) before processing for MS analysis.

#### MS analysis

In-gel digestion as well as LC-MS/MS analysis were performed by the proteomics core facility at EPFL as in the previously published protocol (Grolimund et al., 2013) with minor modifications. In brief each SDS-PAGE gel lane was entirely sliced. Samples were first washed twice for 20 min in 50% ethanol, 50 mM ammonium bicarbonate (AB, Sigma-Aldrich) and dried down by vacuum centrifugation. Sample reduction was then performed with 10 mM dithioerythritol (DTE, Merck-Millipore) at 56°C for one hour. A washing-drying step was performed as described above prior to samples alkylation with 55 mM Iodoacetamide (IAA, Sigma-Aldrich) for 45 min at 37°C light protected. Samples were then washed-dried and stored on ice. Digestion was performed overnight at 37°C using modified Mass Spectrometry grade trypsin (Trypsin Gold, Promega) at a concentration of 12.5 ng/µl in 50 mM AB and 10 mM CaCl2. Resulting peptides were extracted twice for 20 min in 70% ethanol, 5% formic acid (FA, Merck-Millipore) with permanent shaking. Finally, samples were dried down by vacuum centrifugation. Peptides were desalted on C18 StageTips (Rappsilber et al., 2007) and dried down by vacuum centrifugation again. Samples were resuspended in 2% acetonitrile (ACN, Biosolve), 0.1% FA for LC-MS/MS injections. Nano-flow separations were performed on a Dionex Ultimate 3000 RSLC nano UPLC system (Thermo Fischer Scientific) on-line connected with an Orbitrap Elite Mass-Spectrometer (Thermo Fischer Scientific). A home-made capillary pre-column (Magic AQ C18; 3 µm-200 Å; 2cm x 100 µm ID) was used for sample trapping and cleaning. Analytical separation was then performed using a C18 capillary column (Nikkyo Technos Co; Magic AQ C18; 3µm-100Å; 15cm x 75µm ID) at 250 nl/min. Database search was performed using Mascot (Matrix Science), MS Amanda (Dorfer et al., 2014) and SEQUEST in Proteome Discoverer v.1.4 against the Uniprot Human protein database. Strepativin and BirA sequences were manually added. Searches were performed with trypsin cleavage specificity, ion mass tolerance of 10 ppm for the precursor and 0.5 Da for the fragments. Carbamidomethylation was set as a fixed modification, whereas oxidation (M), acetylation (Protein N-term), phosphorylation (STY) were considered as variable modifications.

Raw MS data was analyzed using Scaffold for filtering and to create a non-redundant list of all replicates. Thresholds of 1% protein and peptide level false discovery rate (FDR) and at least two unique peptides per protein were set.

#### Raw data deposition

The mass spectrometry proteomics data have been deposited to the ProteomeXchange Consortium (www.proteomexchange.org) via the PRIDE partner repository with the dataset identifier PXD016984 (**Username:**; **Password:** lnrdY7v7).

### Key Resources Table

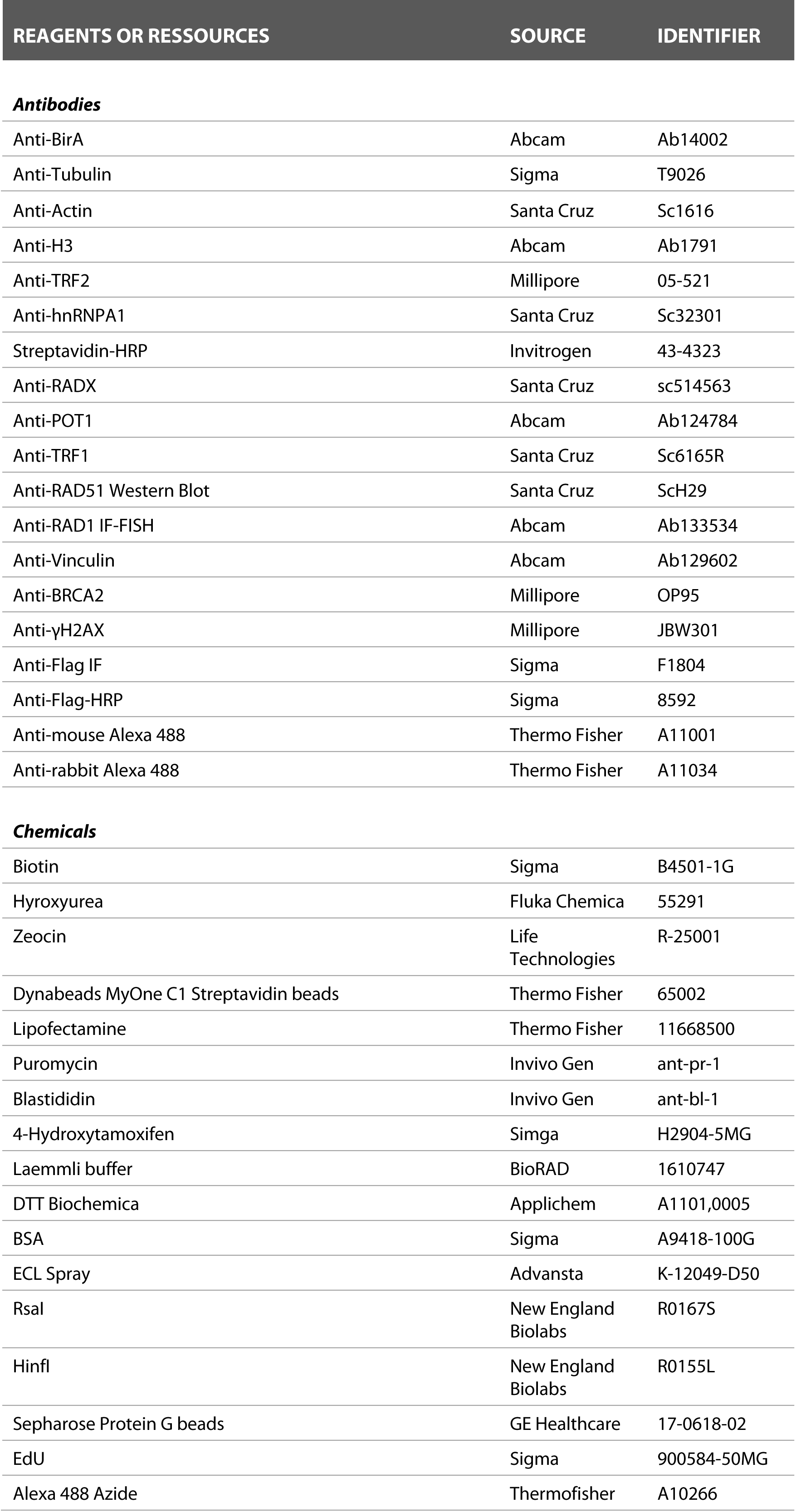
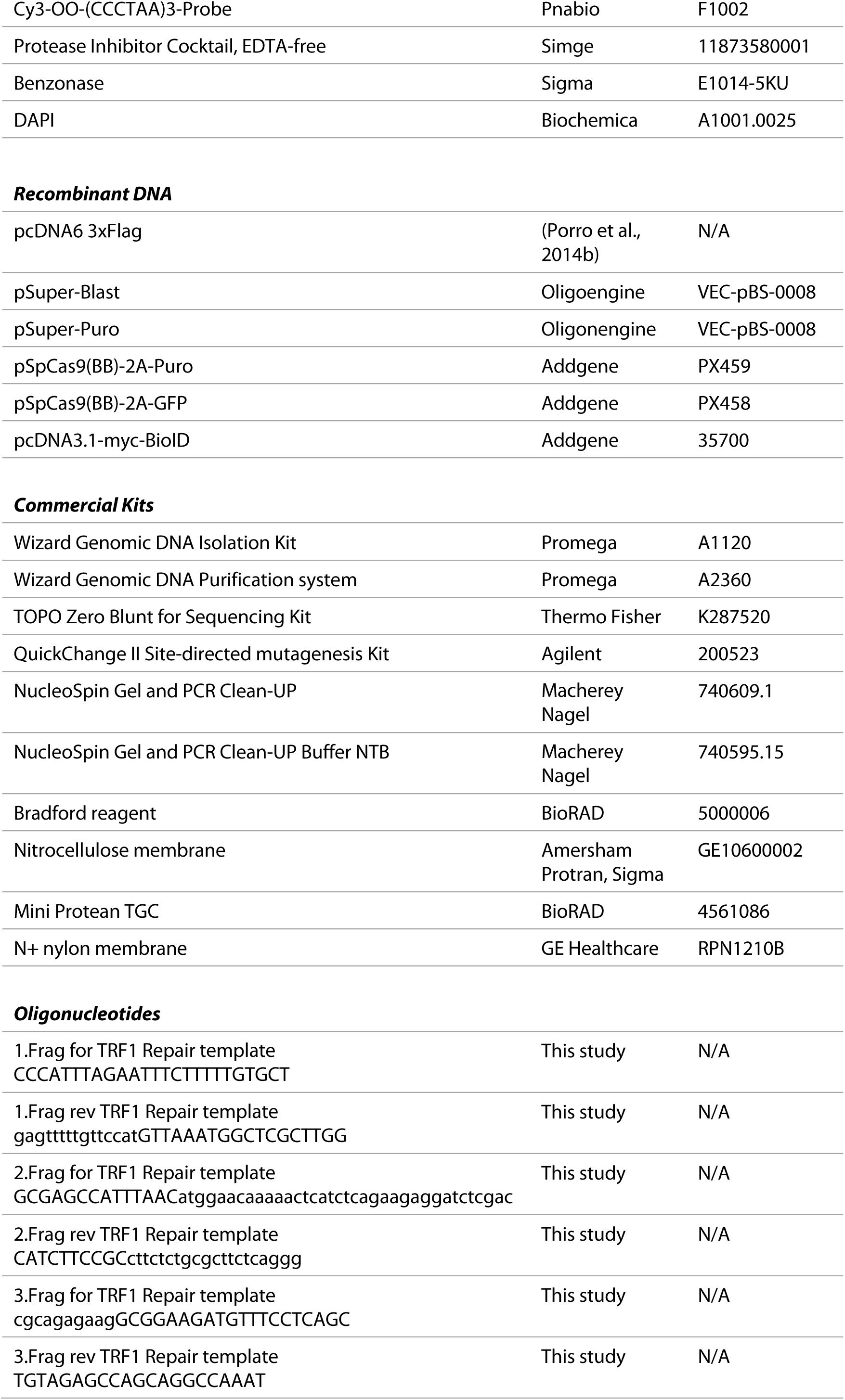
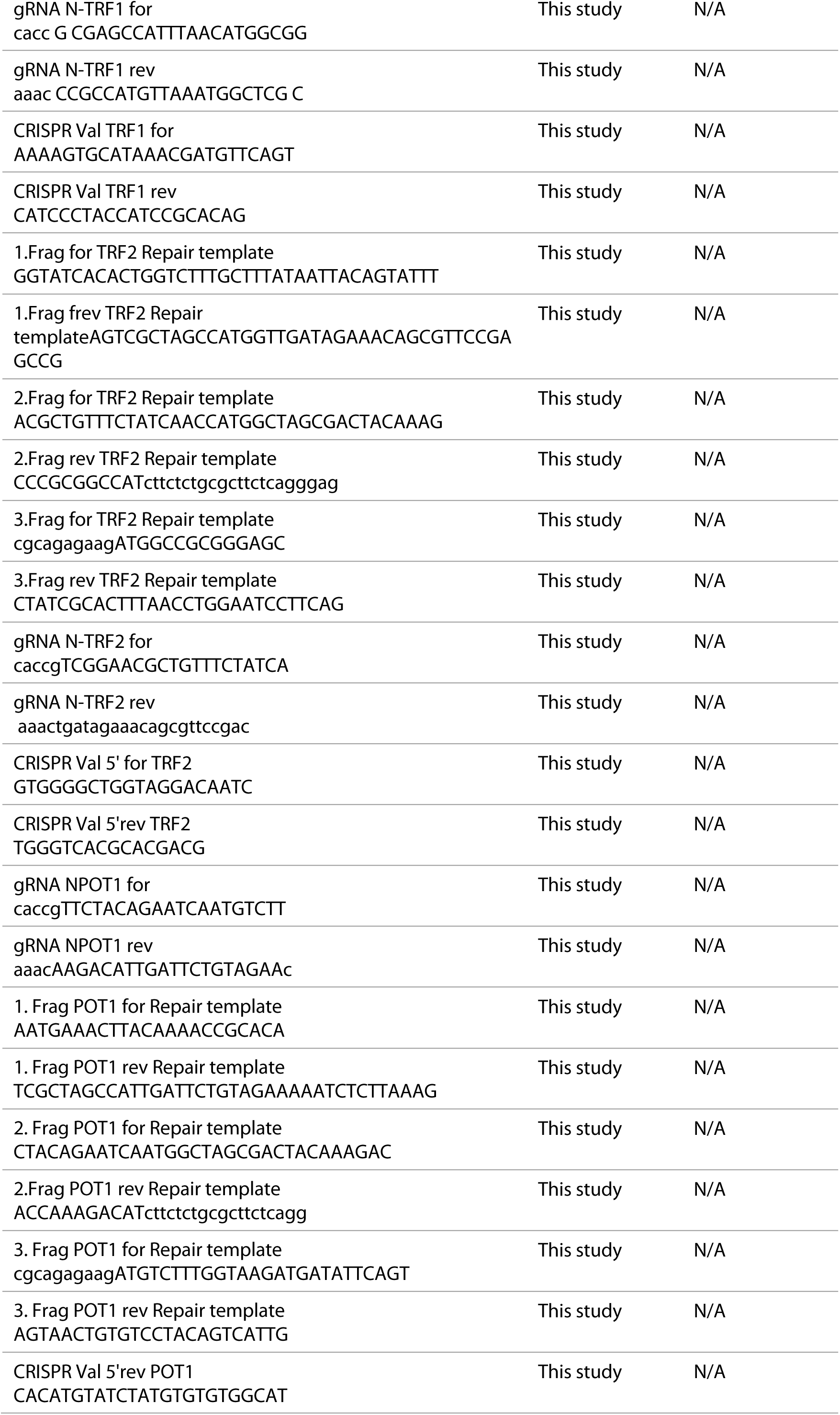
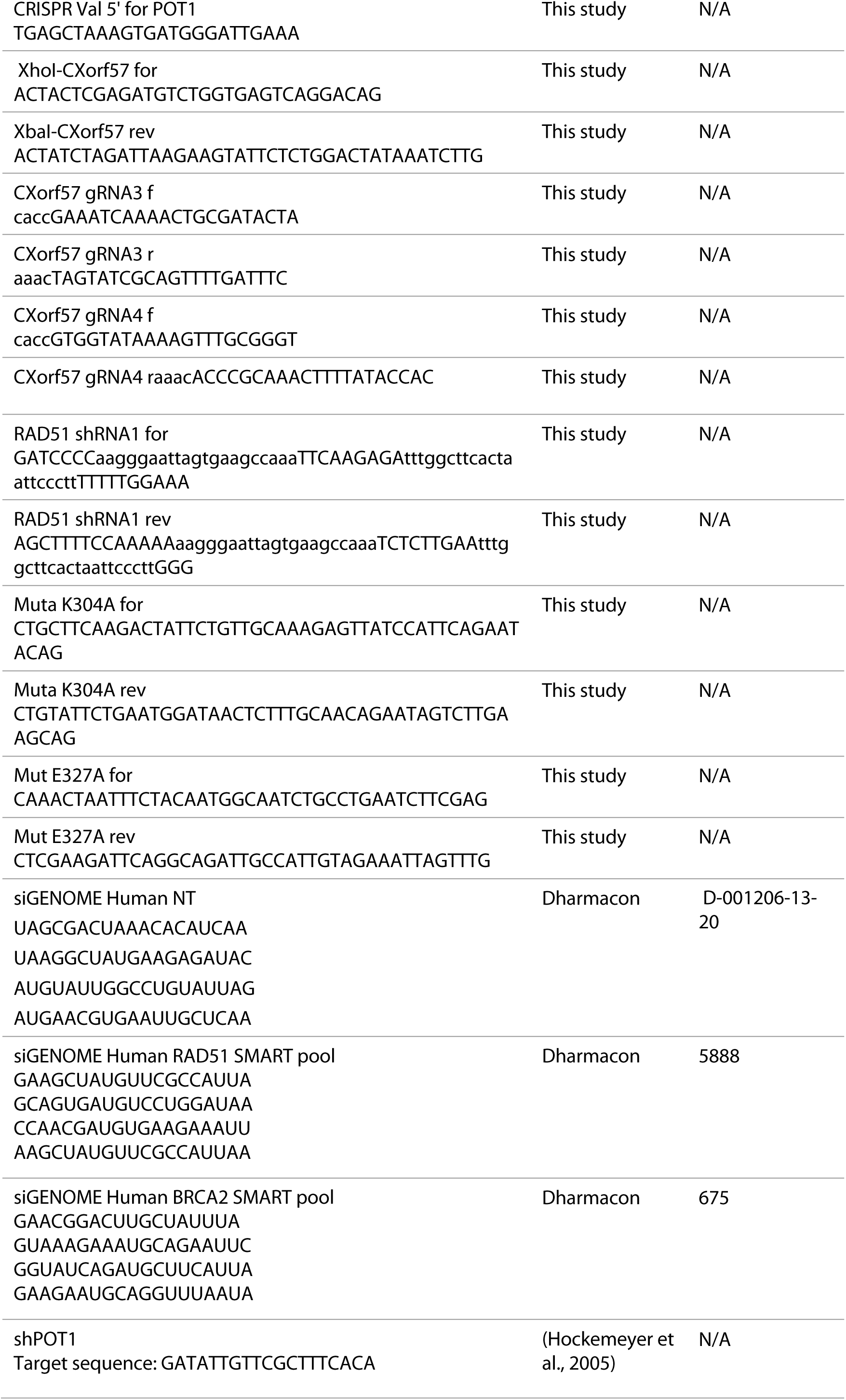
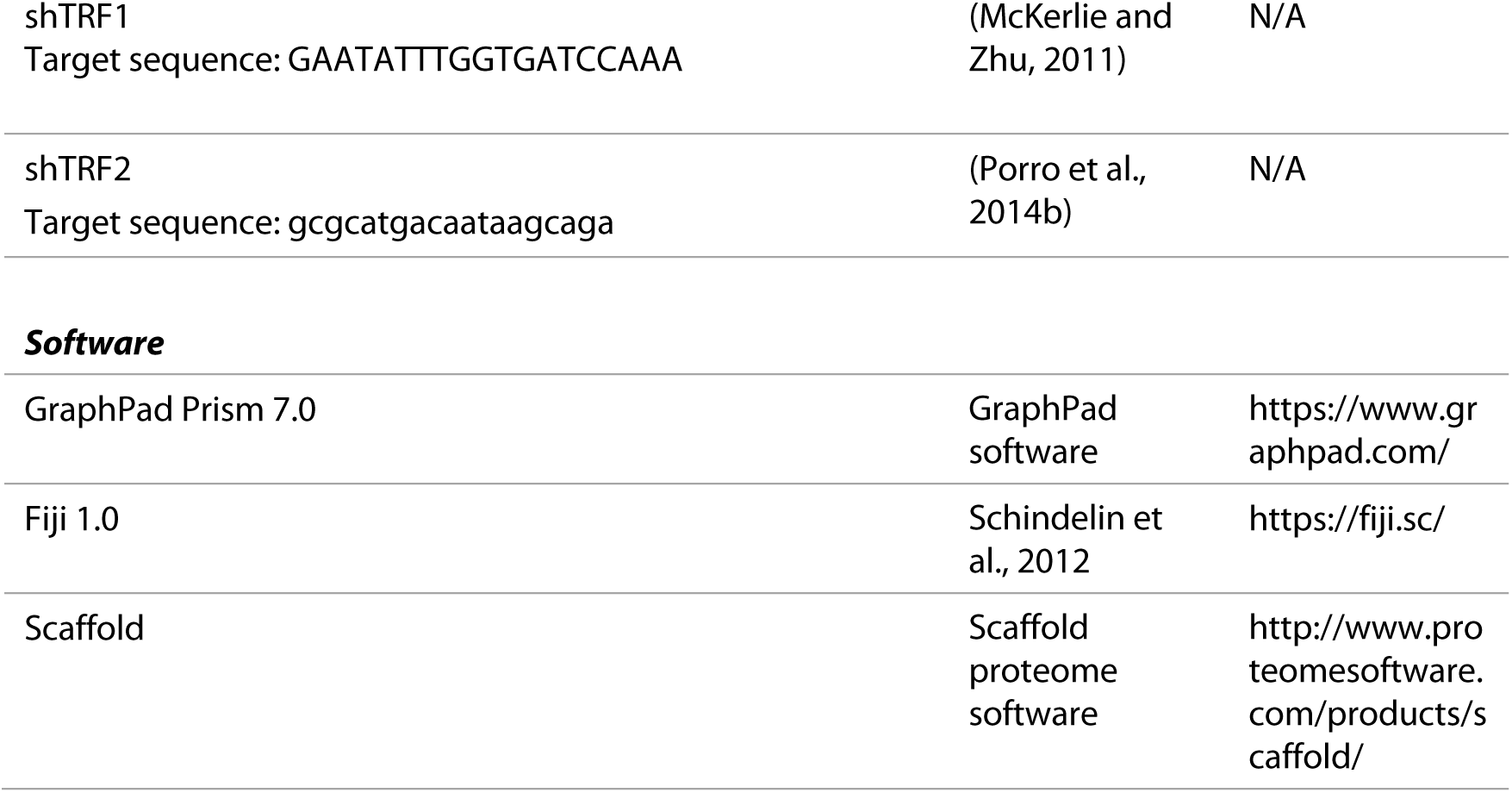

